# Developing an endogenous quorum-sensing based CRISPRi circuit for autonomous and tunable dynamic regulation of multiple targets in industrial *Streptomyces*

**DOI:** 10.1101/2020.01.17.910026

**Authors:** Jinzhong Tian, Gaohua Yang, Yang Gu, Xinqiang Sun, Yinhua Lu, Weihong Jiang

## Abstract

Dynamic regulation has emerged as an effective strategy to improve product titers by balancing metabolic networks, which can be implemented by coupling gene expression to pathway-independent regulatory elements, such as quorum-sensing (QS) systems. However, these QS-based circuits are often created on heterologous systems and must be carefully tuned through tedious testing and optimization process to make them work well, which hampers their application in industrial microbes including streptomycetes. In this study, we design a pathway-independent QS circuit by directly integrating an endogenous QS system with CRISPRi (named EQCi) in the industrial rapamycin-producing strain *Streptomyces rapamycinicus*. EQCi has the advantages of both the QS system and CRISPRi, which enables tunable, fully autonomous and dynamic regulation of multiple targets simultaneously. To demonstrate its effectiveness, we downregulate three key nodes in essential pathways separately to divert metabolic flux toward rapamycin biosynthesis. In each case, significant increases in rapamycin titers are achieved. We further apply EQCi to simultaneously control these three key nodes with proper repression strength by changing sgRNA targeting positions. The final rapamycin titer reaches to 1836±191 mg/L, which is the highest reported titer. Notably, compared to traditional static engineering strategy, which result in growth arrest and suboptimal rapamycin titers, EQCi regulation substantially promote rapamycin titers without affecting cell growth, which indicates that it can achieve the trade-off between essential pathways and product synthesis. Collectively, this study provides a simple and effective strategy for optimizing product titers and may have the potential to apply to other industrial microorganisms.

## INTRODUCTION

Microbes have been engineered as renewable cell factories to efficiently produce a vast array of commercially valuable products, such as pharmaceuticals, biofuels and biochemicals. However, economically viable microbial cell factories are often limited by low product titers and yields. Previously, static metabolic engineering, which normally involves deleting key genes from competitive pathways and overexpressing genes from target pathways by engineering promoters or ribosome binding sites (RBS), are always employed to maximize product titers. But static engineering strategies often result in metabolic imbalance, pathway intermediate accumulation and growth retardation, thereby limiting product titers and yields (*1, 2*). To address this challenge, dynamic metabolic engineering strategies, which could allow microbes to autonomously adjust the metabolic networks in response to environmental changes in real time, have recently been developed and applied for efficient optimization of microbial cell factories (*3-8*).

Dynamic regulation could be mainly implemented in two different manners, namely, pathway-dependent and pathway-independent regulation. Pathway-dependent control systems are generally employed to redirect metabolic fluxes upon sensing changes in concentration of pathway intermediates or byproducts. To execute such dynamic regulation, characterization of a pathway-specific biosensor, such as a metabolite-responsive transcription factor or riboswitch, is a prerequisite (*4, 9*). Using this strategy, significantly improved production of a number of value-added products in various microbes, such as lycopene (*10*), fatty acid ethyl ester (FAEE) (*11*), amorphadiene in *E. coli (12)*, lysine in *Corynebacterium glutamicum(13)*, N-acetylglucosamine (GlcNAc) in *Bacillus subtilis (14)*, have been achieved. Pathway-independent dynamic control, which utilizes common regulatory elements, such as quorum-sensing (QS) circuitry, to couple gene expression to the physiological state of the cell, has also been established and applied to improve product titers and yields (*15-19*). Such dynamic regulation has more broadly applicability and could be implemented across different pathways without developing a specific biosensor. However, to our knowledge, pathway-independent QS circuits are all built on heterologous QS systems, such as the *lux* system from *Vibrio fischeri* (*17, 19*) and the *esa* system from *Pantoea stewartii* (*16-18*). To enable them work well with appropriate switching times and desired regulation strength in hosts, repeated testing, screening and optimization of *cis*-regulatory elements (promoter and RBS) driving the expression of QS circuits, is always required, which hampers their wide application in industrial microbes. To address the above challenges, we aim to develop pathway-independent dynamic control circuits by directly harnessing endogenous QS elements, which are ubiquitous in microbes. In addition, in order to enable efficient regulation of multiple pathways simultaneously, we attempt to integrate the endogenous QS element with the gene repression CRISPRi tool (EQCi) for pathway-independent regulation. CRISPRi, which is developed based on the DNase inactivated CRISPR/Cas endonucleases, such as dCas9 (*20, 21*), has been emerged as effective tool to fine-tune the expression of multiple target genes (*22, 23*). Importantly, using CRISPRi, gene repression strength could be optimized by adjusting the expression of dCas9 and sgRNA, or by changing the targeting position of sgRNA (*21, 24*).

*Streptomyces* is an important industrial microbe that is regarded as the richest known reservoir of antibiotics and other valuable natural products, which have been widely used in medicine, agriculture and animal husbandry. However, in most cases, industrial scale-up production of natural products in *Streptomyces* is limited by their low titers. As natural products biosynthesis compete for common precursors with primary pathways (particularly central carbon metabolism), which are essential for cell growth and cannot be deleted from the genome, we attempt to apply the EQCi strategy to boost natural product titers by dynamically knocking down key nodes in essential pathways. Rapamycin produced by *Streptomyces rapamycinicus* was selected as an example for the validation of our EQCi system. This natural product exhibits a broad range of biological activities, including immunosuppressive, antitumor, neuroprotective/neuroregenerative (*25*). It has been approved by FDA as an immunosuppressant and its derivatives have been approved as antitumor drugs. Recent reports show that rapamycin has anti-aging activity, including significantly extending the life span of yeast, fruit flies, nematodes and mice (*25*), and slowing down the aging of human skin (*26*).

To this end, we construct an EQCi system in *S. rapamycinicus*, in which the *dcas9* gene is under the control of a native QS signal-responsive promoter and sgRNA is under the control of a strong constitutive promoter *j23119*. The design of EQCi, by integrating the endogenous QS element and CRISPRi, can enable us to implement dynamic downregulation of multiple essential pathway genes to redirect metabolic flux toward rapamycin biosynthesis in a cell density-dependent manner. EQCi-mediated regulation was applied to downregulating three respective key nodes in TCA cycle, the fatty acid (FA) and aromatic amino acid (AAA) synthesis pathways and resulted in significantly enhanced rapamycin titers. We further apply EQCi to implement downregulation of three key nodes simultaneously and achieve the highest reported titer through screening a library of sgRNA combinations with varying repression strength. The data clearly demonstrated that EQCi-based regulation is a simple and effective strategy for balancing multiplex essential pathways to promote product titers and can be broadly applied to other industrial microorganisms.

## Results

### Design of an endogenous QS system-based CRISPRi circuit (EQCi) for autonomous and dynamic gene regulation in *S. rapamycinicus*

In *Streptomyces*, the QS system is critical to coordinate the initiation of natural product biosynthesis, which is achieved by γ-butyrolactones (GBLs) and their cognate receptors (always TetR family regulators) (*27*). At a low cell density, the receptor binds to its target DNA sites and represses gene transcription. Upon binding to GBL (at a high cell density), the receptor dissociates from target promoters, resulting in transcription derepression. The QS systems have been intensively investigated in several streptomycetes, such as ArpA/AfsA from *Streptomyces griseus* (*28*), ScbR/ScbA from *Streptomyces coelicolor* (*29, 30*) and SbbR/SbbA from *Streptomyces bingchengensis* (*31*). Among them, AfsA, ScbA and SbbA are the GBL synthases responsible for GBL biosynthesis, while ArpA, ScbR and SbbR function as the GBL receptors. BLASTp analysis revealed that there was a possible GBL system encoded by *M271_07485* and *M271_07490*, named as SrbA and SrbR, in *S. rapamycinicus*. They had high amino acid sequence identities with the GBL synthases and receptors from *S. coelicolor* and *S. bingchengensis* (Fig. S1A). Furthermore, we found that the promoter region of *srbA* showed high DNA sequence identity with that of *sbbA* from *S. bingchengensis* (Fig. S1B), whose transcription is directly regulated by SbbR (*31*). This clearly suggests that the promoter region of *srbA* (*srbA*p) might be under the control of SrbR and *srbA*p is possible a GBL-responsive promoter in *S. rapamycinicus*.

To characterize the activity of *srbA*p, a *srbA*p-driving expression of a thermophilic *lacZ* (*32*) cloned on an integrative plasmid pSET152 was constructed, resulting in the plasmid pSET-*srbA*p-*lacZ*. As a positive control, the recombinant plasmid pSET-*ermEp*\*-*lacZ*, in which the strong constitutive promoter *ermEp*\* was used to drive *lacZ* expression, was also constructed (Fig. 1A). Two plasmids were individually introduced into *S. rapamycinicus* 2001, respectively. As an empty control, 2001 with pSET152 (2001-p) was also constructed. No difference in bacterial growth was observed upon LacZ expression, compared to that of the parental strain 2001 as well as 2001-p (data not shown). Continuous measurements of LacZ enzyme activity revealed that strain with pSET-*ermEp*\*-*lacZ* had a very high enzyme activity even at a low cell population density, which was due to constitutive expression of *ermEp*\*. However, strain harboring pSET-*srbAp*\*-*lacZ* showed high enzyme activity only when cell density reached a high level (Fig. 1B). This phenomenon is consistent with our understanding of the working model of the QS system in *Streptomyces*, clearly indicating that *srbAp* is a GBL-responsive promoter.

**Fig. 1.**
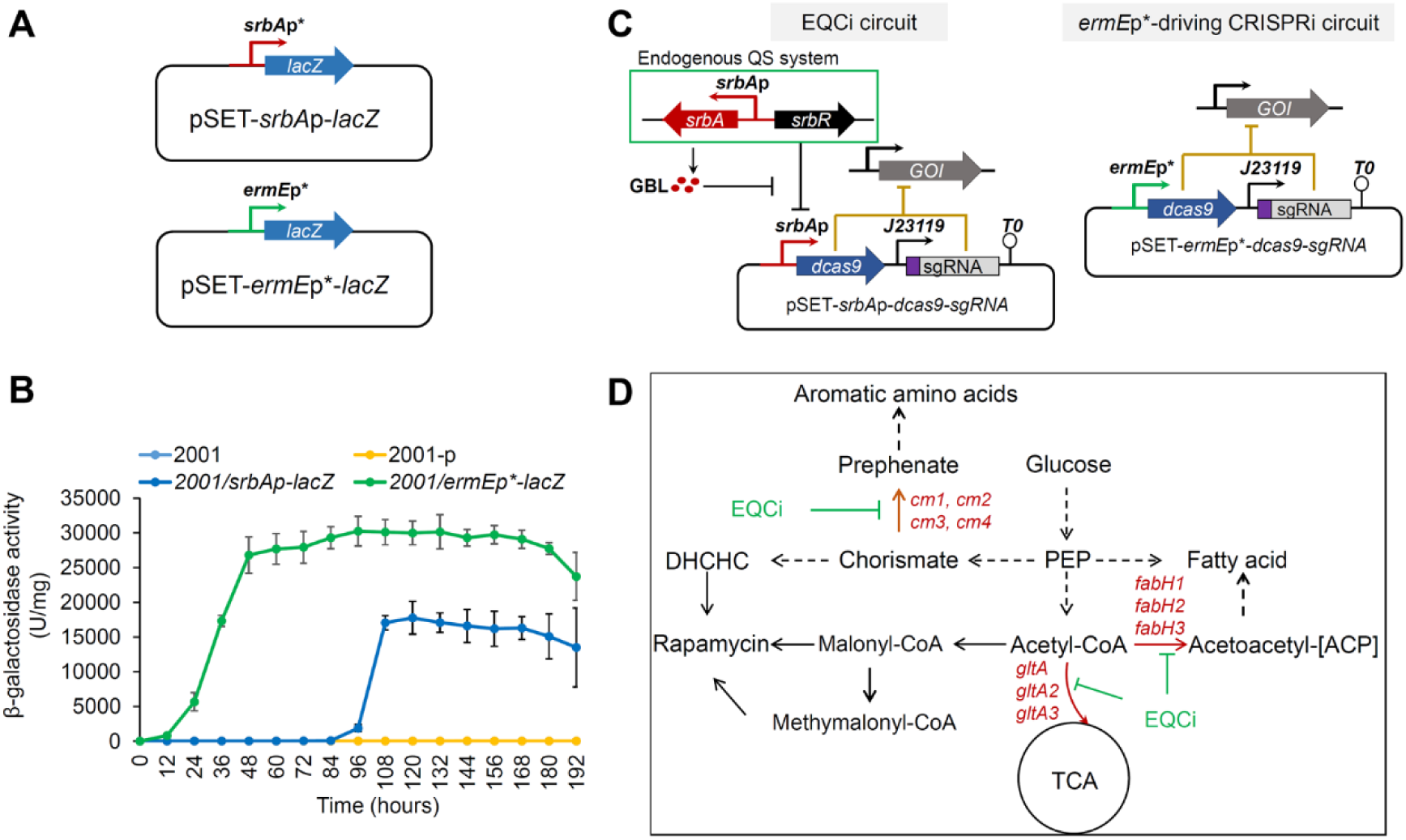
Characterization of the *srbAp* promoter and design of the EQCi genetic circuit in *S. rapamycinicus*. **(A)** Schematic diagram of two reporter systems in which the *lacZ* gene was under the control of *ermEp*\*- and *srbAp*, respectively. The integrative plasmid pSET152 was used as the backbone for the construction of the reporter systems. **(B)** β-galactosidase activities normalized to the biomass of *S. rapamycinicus* harboring the reporter plasmid, pSET-*ermEp*-lacZ* or pSET-*srbAp*-lacZ*. The parental strain 2001 and 2001 with the empty plasmid pSET152 (2001-p) were used as controls. Error bars represent SD of three biological replicates. **(C)** Architecture of the EQCi-based genetic circuit. The integrative plasmid pSET152 was used as the backbone for the construction of EQCi genetic circuits, in which the GBL-responsive promoter *srbA*p was used to drive dCas9 expression and the constitutive strong promoter *j23119* was used to control the transcription of sgRNA targeting the gene of interest (GOI). At a low cell population density, the GBL receptor SrbR binds to target DNA sites in *srbAp*, resulting in the repression of dCas9 expression and then switching off the EQCi circuit. When the cell density is high and the GBL concentration reaches a threshold level, SrbR will dissociate from *srbA*p and lead to dCas9 expression, thereby switching on the EQCi circuit and repressing the transcription of target genes. The *ermEp*\*-driving CRISPRi circuit for static regulation was used as a negative control, in which the constitutive strong promoter *ermEp*\* was used to drive dCas9 expression and *j23119* was used to control sgRNA transcription. **(D)** Schematic diagram of some essential metabolic pathways in *S. rapamycinicus*. Three key nodes selected as the target of EQCi-mediated downregulation are shown by red color. Each node involves 3-4 genes as indicated. EQCi-mediated dynamic repression will divert metabolic flux from essential pathways to rapamycin biosynthesis in a cell population density-dependent manner.

Then, the EQCi circuit was designed, in which *srbA*p was used to drive dCas9 expression and the well-characterized strong promoter *j23119* was used to control the transcription of single guide RNA(s) (sgRNA) targeting the gene of interest (Fig. 1C). At a low cell population density of *S. rapamycinicus* (it means GBL concentration is also low), the GBL receptor SrbR binds to *srbAp*, resulting in the repression of dCas9 expression and then switching off the EQCi circuit. When the cell density is high and the GBL concentration reaches a threshold level, SrbR will no longer bind to the *srbA* promoter, leading to dCas9 expression and thereby switching on EQCi circuit. If it is this case, this EQCi circuit could enable us to dynamically repress the expression of any gene of interest (GOI) or a combination of multiple genes in a cell density-dependent manner and be used to favor accumulating rapamycin by rebalancing metabolic flux distribution between bacterial growth and rapamycin biosynthesis. Here, key nodes from three essential pathways, including TCA cycle, the FA and AAA synthesis pathway (Fig. 1D), were selected to be dynamically downregulated by the EQCi circuits to divert carbon flux away from primary metabolism toward rapamycin biosynthesis, thereby boosting rapamycin titers in *S. rapamycinicus*. For comparison, *ermEp*\*-driving CRISPRi circuits for static downregulation, in which *ermEp\** was used to drive dCas9 expression, were also constructed in some cases (Fig. 1C).

### EQCi-mediated dynamic control of the FA synthesis pathway

Biosynthesis of each rapamycin molecule need 7 malonyl-CoA extension units (*25*). Low endogenous malonyl-CoA levels are always a limiting factor for rapamycin biosynthesis. Here, as a proof of concept, we applied EQCi genetic circuits to elevate malonyl-CoA by dynamically downregulating FA synthesis pathway. As the 3-oxoacyl-ACP synthase is the key enzyme that controls the entry of acetyl-CoA and malonyl-CoA into FA synthesis pathway, genes encoding this key node was chosen for dynamic regulation. In *S. rapamycinicus*, three genes, including *fabH1 (M271_17610), fabH2 (M271_33380*) and *fabH3 (M271_41260)*, were predicted to be responsible for this key node. For each gene, one sgRNA was designed using the online software CRISPRy, targeting the nucleotide (nt) positions of nt 27-46 (*fabH1*) (*sg-fabH1*), nt 8-27 (*fabH2*) (*sg-fabH2*) and nt 179-198 (*fabH3*) (*sg-fabH3*) with respect to the start codons. As controls, *ermEp*\*-driving CRISPRi circuits for static downregulation were also constructed. Totally, six CRISPRi plasmids, namely pSET-*srbAp*-*dcas9*/*sg*-*fabH1*, pSET-*srbAp*-*dcas9*/*sg*-*fabH2*, pSET-*srbAp*-*dcas9*/*sg*-*fabH3*, pSET-*ermEp\**-*dcas9*/*sg*-*fabH1*, pSET-*ermEp\**-*dcas9*/*sg*-*fabH2* and pSET-*ermEp\**-*dcas9*/*sg*-*fabH3*, were constructed and introduced into *S. rapamycinicus* 2001, respectively. Analysis of rapamycin production revealed that EQCi circuit-mediated dynamic repression of *fabH1, fabH2* and *fabH3* all resulted in significantly elevated rapamycin production. The highest titer was achieved by dynamic repression of *fabH3*, which reached 808±21 mg/L, approximately 230% higher than 2001-p*dcas9* (2001 with the plasmid of pSET*dcas9-tracRNA*) or 2001 (Fig. 2A). However, almost no rapamycin production was detected from strains with *ermEp*\*-driving CRISPRi circuits (data not shown). Growth curve analysis revealed that strains with the EQCi circuit for downregulation of *fabH1* and *fabH2* accumulated slightly less biomass than those of 2001 and 2001-*pdcas9*, and strain for dynamic regulation of *fabH3* grew well as that of the controls (Fig. 2B). However, three strains with static regulation circuits (*ermEp*\*-driving dCas9 expression) showed drastically impaired bacterial growth compared to two control strains (Fig. S2A).

**Fig. 2.**
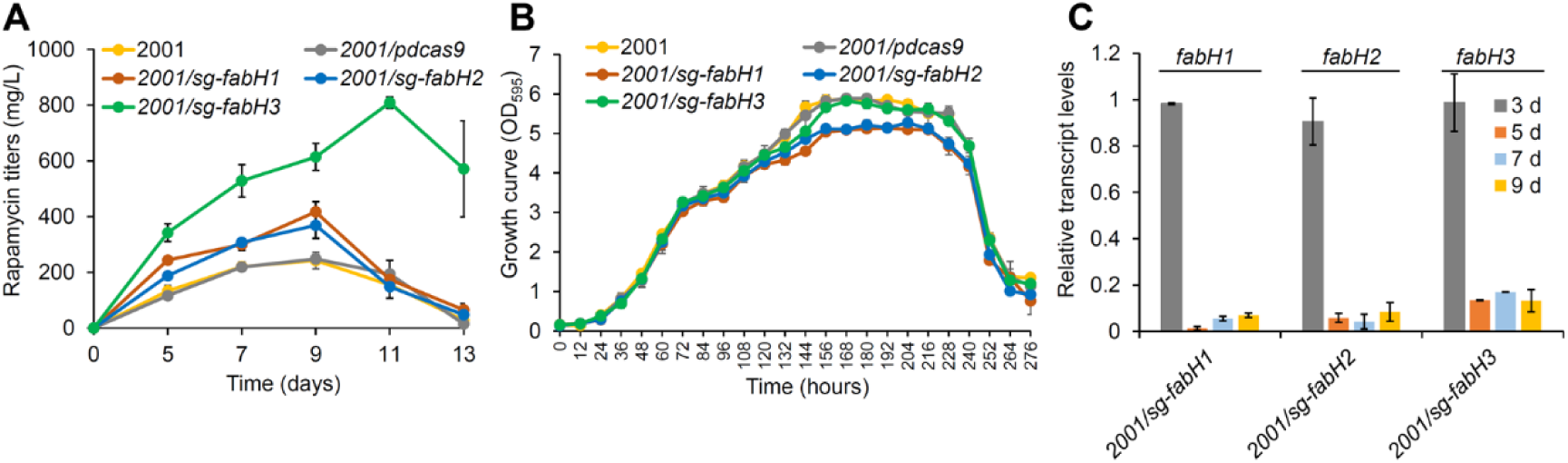
Effects of EQCi-mediated dynamic repression of the key node FabH (3-oxoacyl-ACP synthase) in the FA synthesis pathway. **(A)** Rapamycin titers in strains with the EQCi circuits harboring sgRNA targeting *fabH1-H3* (encoding FabH), respectively. For the construction of the EQCi circuits targeting *fabH1-H3*, one sgRNA (sg) was designed for each gene. Strains with the corresponding CRISPRi circuit are indicated by 2001/*sg-fabH1*, 2001/*sg-fabH2*, and 2001/*sg-fabH3*, respectively. Samples were harvested from *S. rapamycinicus* strains grown in fermentation medium for 5, 7, 9, 11 and 13 days, respectively. The best rapamycin producer is the strain with EQCi circuit harboring the sgRNA targeting *fabH3* (*sg-fabH3*). The parental strain 2001 and 2001 with the recombinant plasmid pSET-*srbAp*-*dcas9* (2001/*pdcas9*) were used as controls. Error bars denote SD of three biological replicates. **(B)** Growth curves of *S. rapamycinicus* strains harboring the EQCi circuits targeting *fabH1-H3*. Samples were harvested from fermentation medium at the time points as indicated and the interval time is 12 hours. Strains 2001 and 2001/p*dcas9* were used as controls. Error bars denote SD of three biological replicates **(C)** Transcriptional analysis of *fabH1-H3* in strains harboring the corresponding EQCi circuit by RT-qPCR. RNA samples were isolated from *S. rapamycinicus* strains, 2001/*sg-fabH1*, 2001/*sg-fabH2* and 2001/*sg-fabH3*, grown in fermentation medium for 3, 5, 7 and 9 days, respectively. *hrdB* (*M271_14880*) was used as the internal control. Error bars represent the standard deviations (SD) from three biological replicates.

To further characterize the behavior of the EQCi circuits, we analyzed the transcript levels of *fabH1, fabH2* and *fabH3* in strains with either the EQCi or *ermEp*\*-driving CRISPRi circuits. The transcript levels of *fabH1, fabH2* and *fabH3* in strains with *ermEp*\*-driving CRISPRi circuits were drastically downregulated to 6.7-14%, 3.3-8.2% and 7.2-12.4% as those of the control strain 2001-*pdcas9* throughout the tested time course (3, 5, 7 and 9 days), respectively (Fig. S2B). However, the EQCi circuits had little effect on the transcription of *fabH1, fabH2* and *fabH3* at early growth stage (3 days); however, from day 5, significant decreases in gene transcription were detected in strains with the EQCi circuits (Fig. 2C), clearly indicating that gene repression is exerted in a cell density-dependent manner. The transcription data obtained here clearly revealed that static, constitutive downregulation of the key node in FA pathway resulted in too early growth arrest and then too low biomass accumulation, which is the reason for suboptimal rapamycin titers. However, the EQCi circuits for dynamic repression was switched on at relatively late growth stage, which could result in build-up of enough biomass and accumulation of high rapamycin titers. Collectively, we have successfully developed a novel pathway-independent dynamic regulation circuit (EQCi), which could enable dynamic regulation of the key node in FA pathway and achieve the balance between of bacterial growth and rapamycin biosynthesis, resulting in significantly elevated rapamycin titers in *S. rapamycinicus*.

### EQCi-mediated regulation of the TCA cycle

Two precursors for rapamycin biosynthesis, malonyl-CoA and methylmalonyl-CoA (*25*), are mainly derived from acetyl-CoA. TCA cycle, which is essential for cell growth, consumes a large number of acetyl-CoA. We aimed to apply the EQCi circuit to dynamically inhibit TCA cycle and divert acetyl-CoA flux toward malonyl-CoA and methylmalonyl-CoA biosynthesis in a cell density-dependent manner, thereby improving rapamycin titers. In TCA cycle, the citric acid synthase catalyzes the formation of citric from acetyl-CoA and oxaloacetic acid, which is the first step of TCA cycle and controls the consuming amount of acetyl-CoA. In the *S. rapamycinicus* genome, there exist three predicted genes (*M271_28125, M271_31465* and *M271_34825*) encoding citrate synthases, which were named as *gltA, gltA1* and *gltA2*, respectively. They were chosen as the target genes for EQCi-mediated dynamic regulation. The sgRNAs of *gltA, gltA1*and *gltA2* were designed to target the positions of nt 137-156 (*sg-gltA*), 8-27 (*sg-gltA1*) and 197-216 (*sg-gltA2*) with respect to the start codon, respectively. Six CRISPRi plasmids (Table S1), including three for dynamic regulation, namely, pSET-*srbAp*-*dcas9/sg*-*gltA*, pSET-*srbAp*-*dcas9*/*sg*-*gltA1* and pSET-*srbAp*-*dcas9*/*sg*-*gltA2*, and three for static regulation, namely, pSET-*ermEp\**-*dcas9*/*sg*-*gltA*, pSET-*ermEp\**-*dcas9*/*sg*-*gltA1* and pSET-*ermEp\**-*dcas9*/*sg*-*gltA2* were constructed and introduced into *S. rapamycinicus*, respectively. Analysis of rapamycin production showed that EQCi-mediated silencing of three respective *gltA* gene resulted in significantly enhanced rapamycin titers. The maximal rapamycin titers of these engineered strains reached approximately 544±66 mg/L (*gltA*), 471±23 mg/L (*gltA1*) and 589±46 mg/L (*gltA2*), approximately 125%, 95% and 143% higher than 2001-p*dcas9* or 2001, respectively (Fig. 3A). However, almost no rapamycin production was detected in strains with the *ermEp*\*-driving CRISPRi circuits (data not shown). Compared to static regulation circuits, which resulted in drastically decreased biomass accumulation (Fig. S3A), the EQCi circuits for individual repression of three *gltA* genes had little or no effect on cell growth (Fig. 3B). We additionally showed that, compared to highly reduced transcription of *gltA, gltA1* and *gltA2* in strains with static regulation circuits throughout the tested time course (3, 5, 7 and 9 days) (Fig. S3B), the EQCi circuits resulted in gene downregulation in a cell density-dependent manner (Fig. 3C), similarly as detected for dynamic regulation of three *fabH* genes. Therefore, it could be concluded that, the EQCi circuits could achieve the coordination of metabolic fluxes between TCA cycle and rapamycin biosynthesis, thereby resulting in significantly elevated rapamycin titers.

**Fig. 3.**
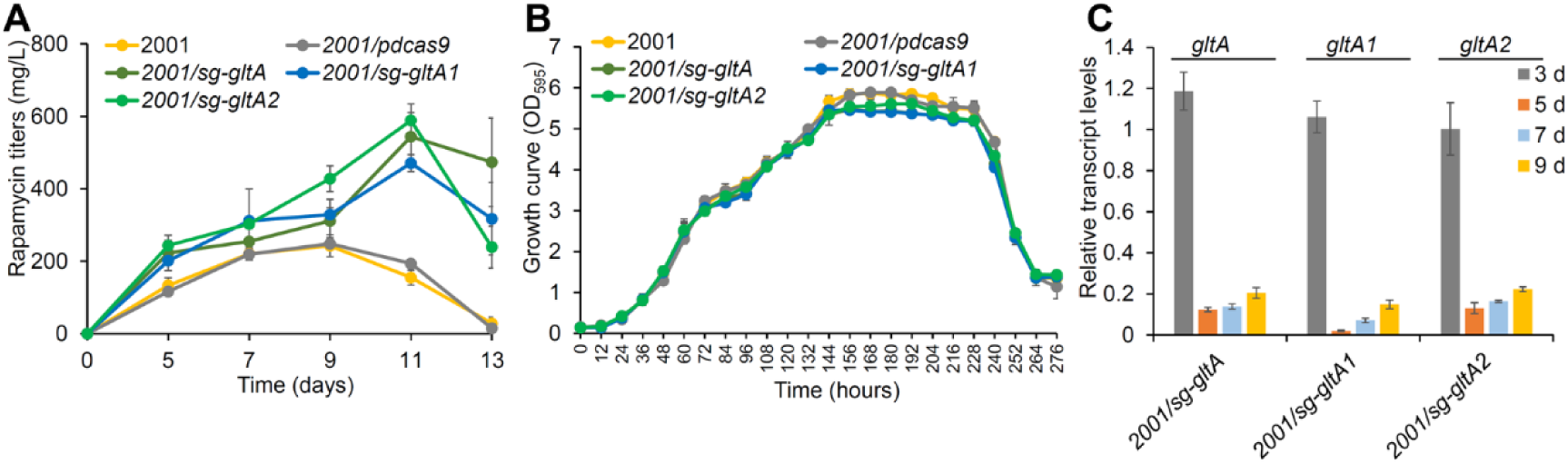
Effects of EQCi-mediated downregulation of the key node (citrate synthase) in TCA cycle. **(A)** Rapamycin titers in strains with the EQCi circuits harboring sgRNA targeting three *gltA* genes (*gltA-A2*, encoding citrate synthase), respectively. For the construction of the EQCi circuits targeting *gltA-A2*, one sgRNA (sg) was designed for each gene. Strains with the corresponding CRISPRi circuit are indicated by 2001*/sg-gltA*, 2001*/sg-gltA1* and 2001/*sg-gltA2*, respectively. Samples were harvested from *S. rapamycinicus* strains grown in fermentation medium for 5, 7, 9, 11 and 13 days, respectively. The best rapamycin producer is the strain with EQCi circuit harboring the sgRNA targeting *gltA2*. Strains 2001 and 2001 with the recombinant plasmid pSET-*srbAp*-*dcas9* (2001/*pdcas9*) were used as controls. Error bars denote SD of three biological replicates. **(B)** Growth curves of *S. rapamycinicus* strains harboring the EQCi circuits targeting *gltA-A2*. Samples were harvested from fermentation medium at the time points as indicated and the interval time is 12 hours. Strains 2001 and 2001/p*dcas9* were used as controls. Error bars denote SD of three biological replicates **(C)** Transcriptional analysis of *gltA-A2* in strains harboring the corresponding EQCi circuit by RT-qPCR. RNA samples were isolated from *S. rapamycinicus* strains, 2001/*sg-gltA*, 2001/*sg-gltA1* and 2001/*sg-gltA2*, grown in fermentation medium for 3, 5, 7 and 9 days, respectively. *hrdB* was used as the internal control. Error bars represent the standard deviations (SD) from three biological replicates.

### EQCi-mediated downregulation of the AAA synthesis pathway

DHCHC is the starter unit of rapamycin synthesis pathway (*25, 33*). It is derived from the hydrolysis of chorismate that is naturally required for aromatic amino acids (AAA) synthesis, including phenylalanine (Phe), tyrosine (Tyr) and tryptophan (Trp). Given that AAAs are essential amino acids, we applied the EQCi circuits to dynamically control chorismate flux toward AAA biosynthesis, thereby increasing rapamycin titers. In Phe and Tyr biosynthetic pathway, the first step consuming chorismate is catalyzed by chorismate mutase (CM) to form prephenate, which are encoded by four predicted genes, including *M271_28280* (*cm1*), *M271_36305* (*cm2*), *M271_37500* (*cm3*) and *M271_43345* (*cm4*). Therefore, these four genes were chosen as the target genes for dynamic control. The sgRNA of *cm1 to cm4* were designed to target the positions of nt 32-51 (*sg1-cm1*), nt 149-168 (*sg1-cm2*), nt 51-70 (*sg1-cm3*) and nt 22-41 (*sg-cm4*) with respect to the start codons, respectively. Accordingly, four CRISPRi plasmids for dynamic regulation (Table S1), namely, pSET-*srbAp*-*dcas9*/*sg*-*cm1*, pSET-*srbAp*-*dcas9*/*sg*-*cm2*, pSET-*srbAp*-*dcas9*/*sg*-*cm3* and pSET-*srbAp*-*dcas9*/*sg*-*cm4*, were constructed. As controls, four CRISPRi plasmids for static regulation, namely, pSET-*ermEp\**-*dcas9*/*sg*-*cm1*, pSET-*ermEp\**-*dcas9*/*sg*-*cm2*, pSET-*ermEp\**-*dcas9*/*sg*-*cm3* and pSET-*ermEp*-dcas9*/*sg*-*cm4*, were also generated (Table S1). Eight CRISPRi plasmids were individually introduced into *S. rapamycinicus* 2001. Analysis of rapamycin production indicated that all four tested EQCi circuits resulted in significantly elevated rapamycin titers. The maximal titers of four engineered strains reached approximately 316±14 mg/L (*cm1*), 735±52 mg/L (*cm2*), 693±84 mg/L (*cm3*), and 694±31 mg/L (*cm4*), respectively (Fig. 4A). But almost no rapamycin titer were detected in strains with static regulation circuits. We showed that, when the EQCi circuits were employed, except for strain with the EQCi circuit pSET-*srbAp*-*dcas9*/*sg*-*cm1* (repressing *cm1* expression) exhibited a slightly growth arrest, other three strains showed little or no growth changes compared to the controls (2001 and 2001-*pdcas9*) (Fig. 4B). However, static downregulation of these four genes resulted in much less biomass accumulation (Fig. S4A). We further confirmed that transcript levels of these four target genes in strains with dynamic regulation circuits were downregulated in a cell density-dependent manner as described for dynamic regulation of *fabH* and *gltA* genes, while gene transcription was constitutively repressed by static regulation circuits (Fig. 4C). It is worth to note that *cm1* transcription was the most significantly repressed, which may account for the growth arrest as well as lower rapamycin titers of the engineered strain with the dynamic CRISPRi circuit pSET-*srbAp*-*dcas9*/*sg*-*cm1*.

**Fig. 4.**
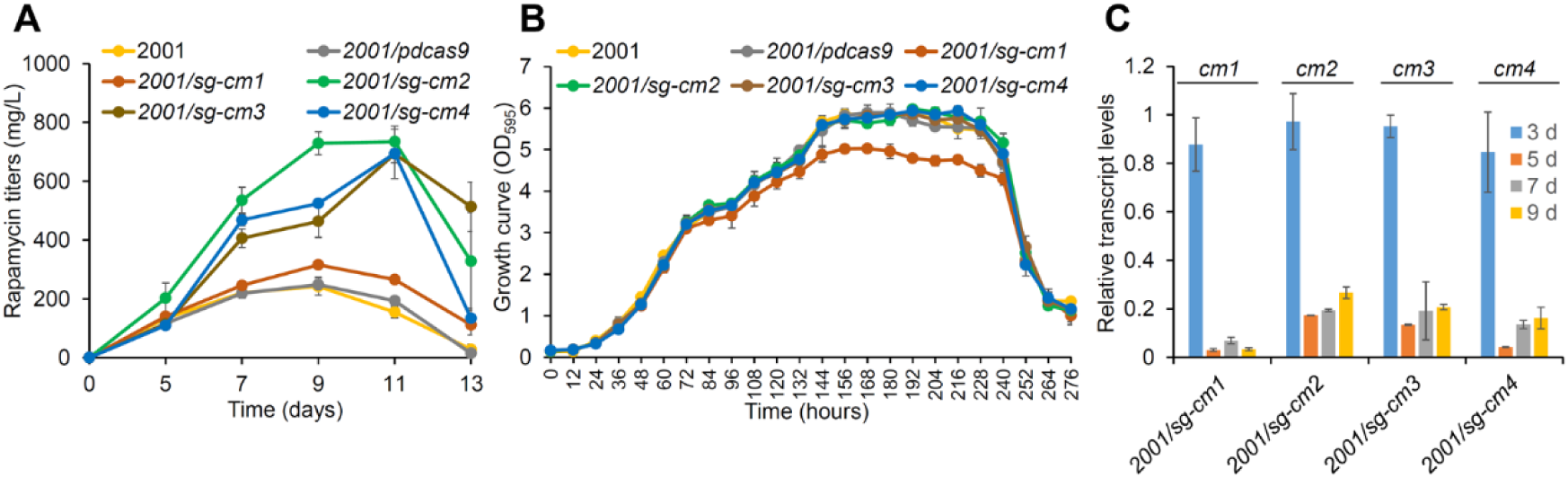
Effects of EQCi-mediated repression of the key node (chorismate mutase) in the AAA synthesis pathway. **(A)** Rapamycin titers in strains with the EQCi circuits targeting four *cm* genes (*cm1-4*, encoding chorismate mutase), respectively. For the construction of the EQCi circuits targeting *cm1-4*, one sgRNA (sg) was designed for each gene. Strains with the corresponding EQCi circuit was indicated by 2001/*sg-cm1*, 2001/*sg-cm2*, 2001/*sg-cm3* and 2001/*sg-cm4*. Samples were harvested from *S. rapamycinicus* strains grown in fermentation medium for 5, 7, 9, 11 and 13 days, respectively. The best rapamycin producer is 2001/*sg-cm2* with EQCi circuit harboring the sgRNA targeting *cm2* (*sg-cm2*). Strains 2001 and 2001 with the recombinant plasmid pSET-*srbAp*-*dcas9* (2001/*pdcas9*) were used as controls. Error bars denote SD of three biological replicates. **(B)** Growth curves of *S. rapamycinicus* strains harboring the EQCi circuits targeting *cm1-4*. Samples were harvested from fermentation medium at the time points as indicated and the interval time is 12 hours. Strains 2001 and 2001/*pdcas9* were used as controls. Error bars denote SD of three biological replicates. **(C)** Transcriptional analysis of *cm1-4* in strains harboring the corresponding EQCi circuit by RT-qPCR. RNA samples were isolated from S. *rapamycinicus* strains, including 2001/*sg-cm1*, 2001/*sg-cm2*, 2001/*sg-cm3* and 2001/*sg-cm4*, grown in fermentation medium for 3, 5, 7 and 9 days, respectively. *hrdB* was used as the internal control. Error bars represent the standard deviations (SD) from three biological replicates.

### EQCi-mediated combinatorial optimization of metabolic networks to substantially increase rapamycin production

As presented above, three key nodes from essential pathways, including TCA cycle, and the FA and AAA synthesis pathways, were selected for dynamic downregulation and achieved significantly enhanced rapamycin titers, respectively. For each key node, which involves 3-4 predicted genes, dynamic regulation of *fabH3, cm2* and *gltA2* resulted in the most significantly improved rapamycin production. Here, we attempted to further increase rapamycin titer by dynamic downregulating these three genes simultaneously using the EQCi circuits. To this end, the EQCi genetic circuit with three sgRNAs targeting *fabH3 (sg-fabH3), gltA2* (*sg-gltA2*) and *cm2* (*sg-cm2*) was constructed and introduced into *S. rapamycinicus*. However, to our surprise, compared to the controls, 2001 and 2001-p*dcas9*, the engineered strain with the circuit targeting three sgRNA simultaneously showed severe growth arrest and decreased rapamycin titers in a cell density-dependent manner (Fig. 5). Before 3-4 days, when the cell density is relatively low, cell growth and rapamycin titers of the engineered strain were comparable with the controls; however, after that, strain with the EQCi genetic circuit accumulated much less biomass and rapamycin titers than the controls. This phenomenon clearly suggests that simultaneous silencing these three target genes at the current regulation strength diverted too much metabolic flux from these three pathways, accordingly resulting in insufficient metabolites supply for primary metabolism and thereby leading to growth arrest and lower rapamycin biosynthesis.

**Fig. 5.**
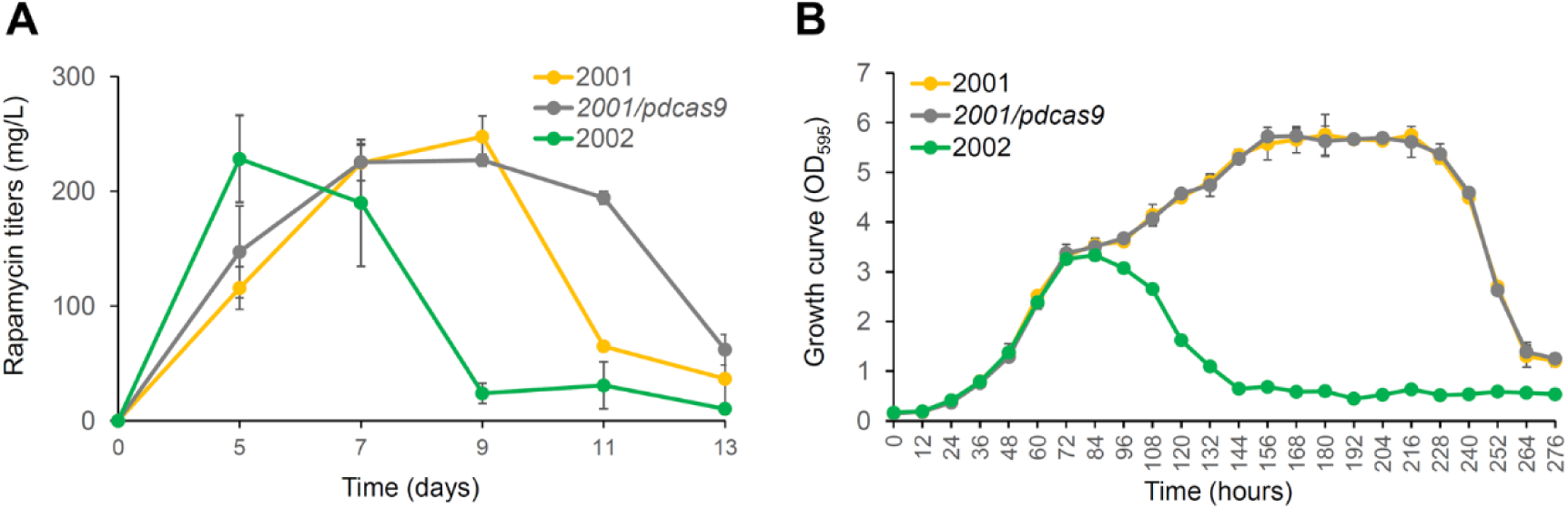
Effects of EQCi-mediated simultaneous downregulation of three key nodes in essential pathways. **(A)** Rapamycin titers in strain with the EQCi circuit-mediated simultaneous repression of *fabH3, gltA2* and *cm2*. Strain with simultaneous repression of *fabH3, gltA2* and *cm2* was indicated by 2002 (containing the EQCi plasmid pSET-*srbAp*-*dcas9-sg-fabH3/*/sg-*gltA2/sg-cm2*). Samples were harvested from *S. rapamycinicus* strains grown in fermentation medium for 5, 7, 9, 11 and 13 days, respectively. 2001 and 2001 with the recombinant plasmid pSET-*srbAp*-*dcas9* (2001/*pdcas9*) were used as controls. Error bars denote SD of three biological replicates. **(B)** Growth curves of strain with EQCi-mediated simultaneous repression of *fabH3, gltA2* and *cm2*. Samples were harvested from fermentation medium at the time points as indicated and the interval time is 12 hours. 2001 and 2001/*pdcas9* were used as controls. Error bars denote SD of three biological replicates.

To search for proper sgRNA combination with desired regulation strength, we sought to create a library of EQCi circuits by changing the combination of three sgRNAs targeting *fabH3, gltA2* and *cm2* with different repression strength. To this end, as the first step, we designed three additional sgRNAs for both *fabH3* and *gltA2*, and two additional sgRNAs for *cm2* (Fig. 6A), and attempted to obtain a series of sgRNAs with different regulation strength. The corresponding EQCi circuits were constructed and introduced in *S. rapamycinicus* 2001. Then, 3-4 engineered strains containing different EQCi circuits for dynamic downregulation of each gene were compared for cell growth, rapamycin titers and gene transcription. We showed that, for each tested gene, obviously different effects were detected by varying the targeting positions of sgRNAs (Fig. 6 and Fig. S5). For dynamic regulation of *fabH3* and *cm2, sg-fabH3* and *sg-cm2* achieved the highest rapamycin titers (Fig. 6B and 6D). For *gltA2, sg2-gltA2* resulted in the maximum rapamycin titer of 676±35 mg/L, approximately 27% higher than that of *sg-gltA2* (Fig. 6C). Transcription analysis confirmed that EQCi-mediated repression is closely associated with the targeting positions of sgRNA (Fig. S5). The closer of sgRNA targeting positions to the start codon, the stronger inhibition effect could be obtained. Growth curve further showed that EQCi circuits harboring the last three sgRNAs for *fabH3* and *gltA2*, or the last two sgRNAs for *cm2* has little effect on bacterial growth (Fig. S5), suggesting they could achieve the trade-offs between bacterial growth and product synthesis. It is worth to note that, compared to *sg-gltA2*, which resulted in drastic downregulation of *gltA2* transcription and obvious less biomass accumulation, *sg2-gltA2* has little effect on cell growth and repressed gene transcription to a less extent (Fig. S5B and E), suggesting a more balanced state between TCA cycle and rapamycin biosynthesis. In addition, EQCi circuits containing sgRNAs targeting too close to the start codon, including *sg1-fabH3, sg1-gltA2* and *sg1-cm2*, resulted in impaired cell growth, decreased rapamycin titers and drastic gene downregulation in a cell density-dependent manner (data not shown). The growth arrest and very lower rapamycin titers of these three engineered strains are clearly owing to drastic downregulation of target genes when the cell density reached a threshold.

**Fig. 6.**
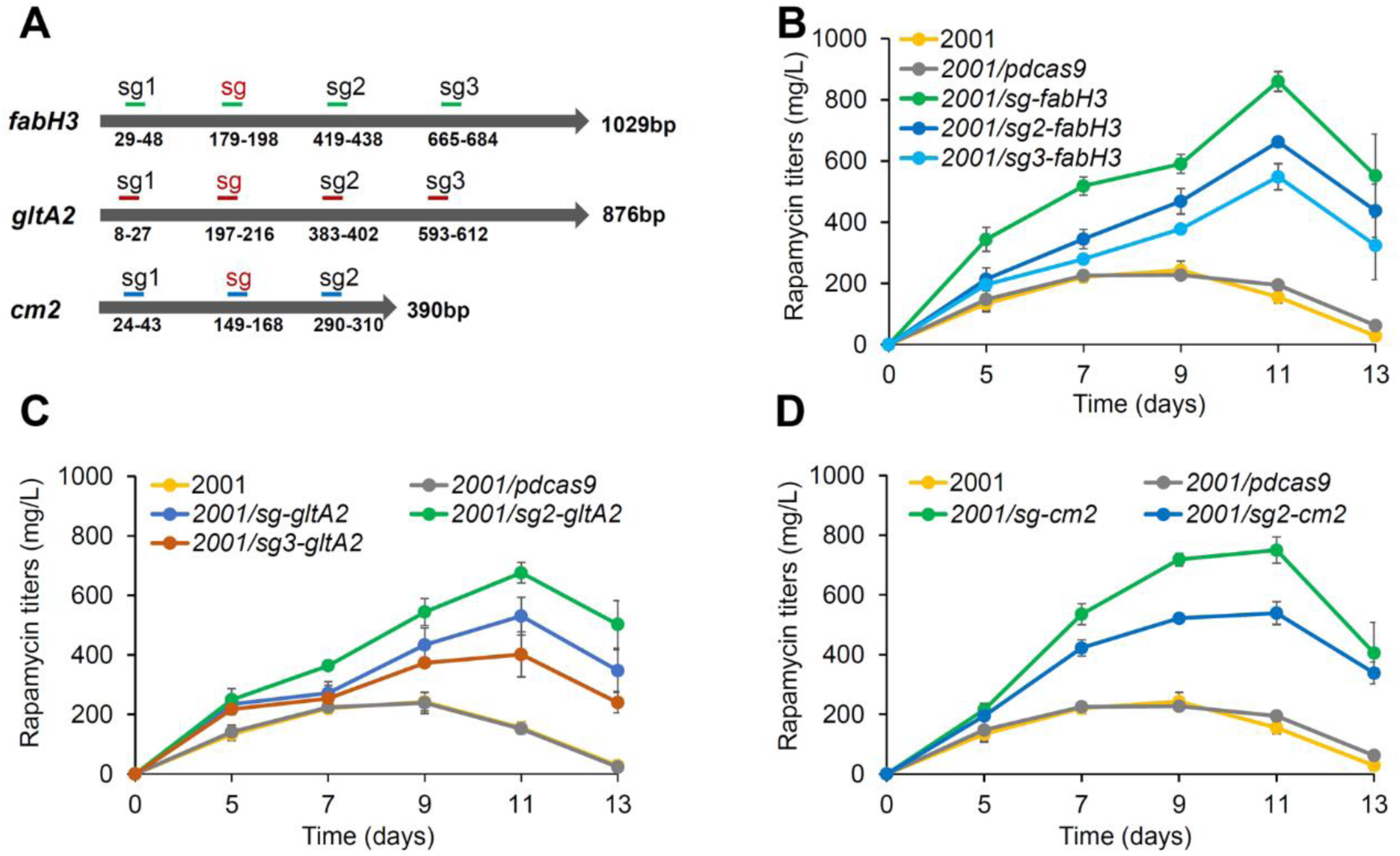
Effects of EQCi-mediated dynamic regulation of three key genes *fabH3, gltA2* and *cm2* on rapamycin titers by varying sgRNA targeting positions. **(A)** Schematic diagram of sgRNA targeting positions designed on the non-template strands of *fabH3, gltA2* and *cm2*. The positions are as indicated. sgRNAs, which have already been tested in above experiments are presented by red font. **(B-D)** Rapamycin titers in strains with the EQCi circuits targeting *fabH3* (three designed sgRNAs), *gltA2* (three designed sgRNAs) and *cm2* (two designed sgRNAs), respectively. Strains with the corresponding EQCi circuit were as indicated. Samples were collected from *S. rapamycinicus* strains grown in fermentation medium for 5, 7, 9, 11 and 13 days, respectively. 2001 and 2001 with the recombinant plasmid pSET-*srbAp*-*dcas9* (2001/*pdcas9*) were used as controls. Error bars denote SD of three biological replicates.

Given that *sg1-fabH3, sg1-gltA2* and *sg1-cm1* resulted in severe growth arrest, they were omitted in library construction. Therefore, totally, 18 EQCi circuits with each combination of three different sgRNAs (3 sgRNAs for both *fabH3* and *gltA2*, two sgRNAs for *cm1*, 3×3×2=18) were constructed and introduced into *S. rapamycinicus* 2001 (Fig. S6). We showed that, among the engineered strains (totally 18 strains) harboring the corresponding EQCi circuit with different sgRNA combination, four strains (2012, 2013, 2016, 2019) produced maximal rapamycin titers of over 1000 mg/L, much higher than those of the controls (247±18 mg/L of 2001 and 239±35 mg/L of 2001-p*dcas9*) (Fig. 7A and Fig. S7). Among them, strain 2013 with the EQCi circuit harboring the sgRNA combination of *sg2-gltA/sg3-fabH3/sg2-cm2* showed the highest rapamycin titer of 1836±191 mg/L, approximately 660% higher than the controls (Fig. 7A). Then, we evaluated growth curves as well as target gene transcription of the top three high-yield rapamycin-producing strains (2012, 2013 and 2019). Little or no difference in cell growth between the engineered strains and the controls was observed (Fig. 7B). We further confirmed that target genes were dynamic regulated in a cell density-dependent manner. Owing to the sgRNA targeting positions located away from the start codons, three target genes were repressed to a less extent from day 5, and the transcript levels were downregulated to approximately 27-57% as those of the control (Fig. 7C-E). The data presented clearly suggest that EQCi-mediated repression of these three key genes at the proper repression strength, particularly the sgRNA combination of *sg2-gltA/sg3-fabH3/sg2-cm2*, were able to balance the metabolic fluxes between growth and rapamycin production.

**Fig. 7.**
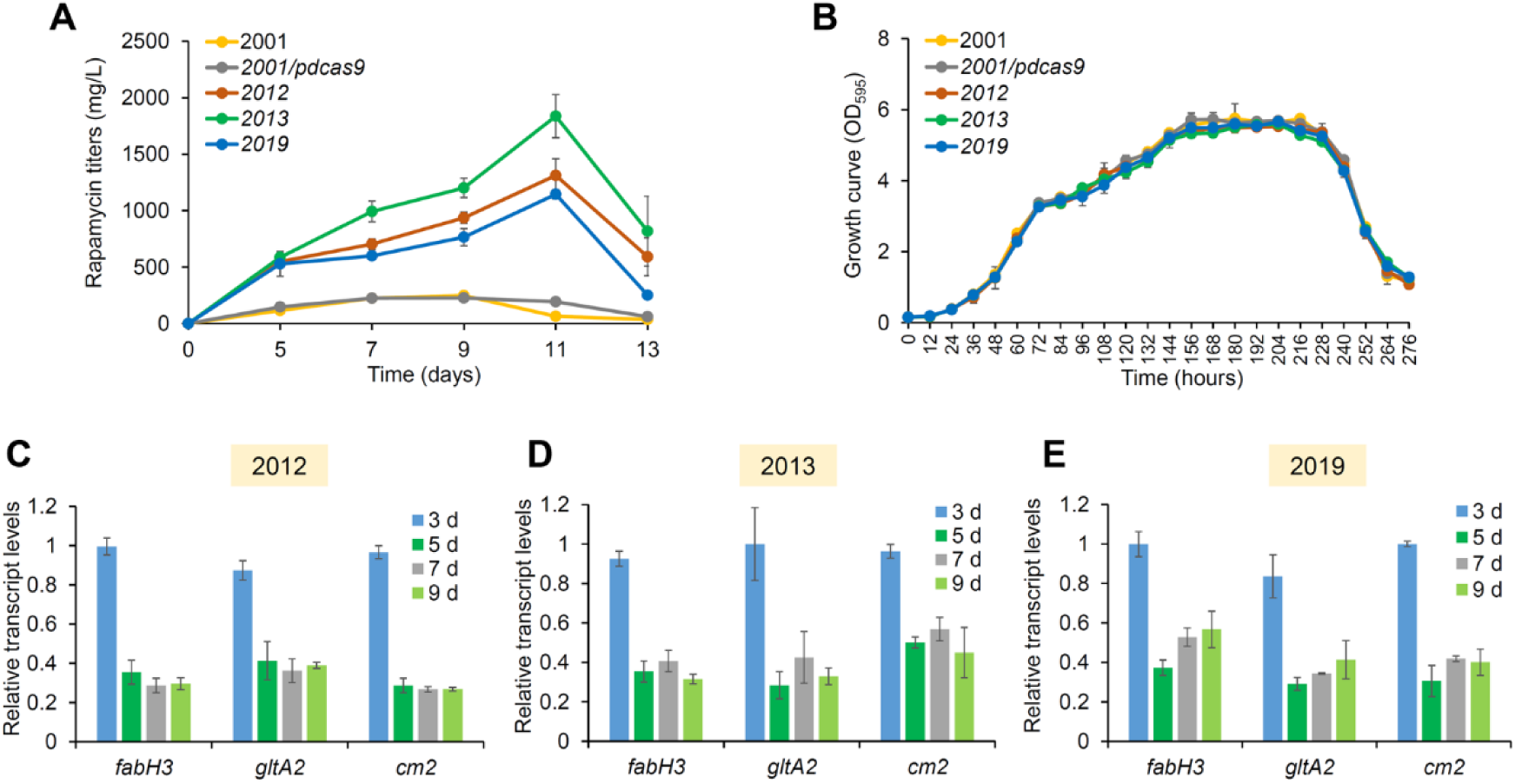
Effects of the EQCi circuits for simultaneous downregulation of three key nodes with proper repression strength. **(A)** Rapamycin titers in strains with EQCi circuit-mediated simultaneous repression of *fabH3, gltA2* and *cm2* with proper repression strength. The top three high-yield rapamycin-producing strains were indicated as 2012 (harboring the EQCi plasmid pSET*-srbAp-dcas9*-*sg2-fabH3/sg3-gltA2/sg-cm2*) (1312±148 mg/L), 2012 (harboring the EQCi plasmid pSET*-srbAp-dcas9*-*sg3-fabH3/sg3-gltA2/sg2-cm2*) (1836±191 mg/L) and 2019 (harboring the EQCi plasmid pSET*-srbAp-dcas9*-*sg3-fabH3/sg3-gltA2/sg2-cm2*) (1144±41 mg/L), respectively. Samples were collected from *S. rapamycinicus* strains grown in fermentation medium for 5, 7, 9, 11 and 13 days, respectively. 2001 and 2001 with the recombinant plasmid pSET-*srbAp*-*dcas9* (2001/*pdcas9*) were used as controls. Error bars denote SD of three biological replicates. **(B)** Growth curves of the top three high-yield rapamycin-producing strains harboring EQCi circuit-mediated simultaneous repression of *fabH3, gltA2* and *cm2* with propter repression strength. Samples were harvested from fermentation medium at the time points as indicated and the interval time is 12 hours. 2001 and 2001*/pdcas9* were used as controls. Error bars denote SD of three biological replicates. **(C-E)** Transcript levels of *fabH3, gltA2* and *cm2* in the top three high-yield rapamycin-producing strains (2012, 2013 and 2019). RNA samples were isolated from S. *rapamycinicus* strains grown in fermentation medium for 3, 5, 7 and 9 days, respectively. *hrdB* was used as the internal control. Error bars represent SD from three biological replicates.

## Discussion

In this study, a novel pathway-independent dynamic control strategy EQCi by integrating an endogenous QS system with the robust CRISPRi tool, was designed and developed in the important industrial strain, *S. rapamycinicus*. The EQCi genetic circuit was successfully applied to implementing fully autonomous, tunable and dynamic downregulation of three key nodes in essential pathways separately or simultaneously to divert metabolic fluxes toward rapamycin biosynthesis in a cell population density-dependent manner, which achieved substantial increases in rapamycin titers. Remarkably, the EQCi-mediated multiplexed control of essential pathway at proper repression strength imposed little or no metabolic burden on cell growth, which indicates it could well coordinate the distribution of metabolic fluxes between growth and rapamycin biosynthesis. Previously, pathway-independent QS circuits have been proven to be effective for improving the titers of target products (*15-19*). However, till now, all of the pathway-independent QS circuits are built on heterologous QS systems. To ensure the effectiveness of QS-based dynamic circuits, repeated screening and optimization of *cis*-regulatory elements (including promoter and RBS) driving the expression of QS modules, involving the construction of promoter-RBS library, are generally required. The process is labor-intensive and time-consuming. The EQCi strategy developed here is very simple and convenient by directly placing dCas9 expression under the control of a QS signal (such as GBL)-responsive promoter, which could easily address the above challenges. Moreover, desired repression efficiency in EQCi regulation could be easily achieved by changing sgRNA targeting positions. Finally, considering that QS systems are widely distributed in microbes as well as the simplicity and specificity of CRISPRi, it could be therefore concluded that EQCi-mediated regulation strategy may be easily expanded to engineer other microbial cell factories.

The EQCi circuit has the advantages of both the QS system and CRISPRi, which enables tunable, fully autonomous and dynamic downregulation of single or multiple pathways to redirect metabolic fluxes, thereby maximizing target product biosynthesis without the need of human intervention of fermentations. Regretfully, in this work, dual-regulation genetic circuits to restrict metabolic flux through primary metabolism and siphon flux into rapamycin biosynthesis simultaneously was not carried out. It could be imagined that if dynamic upregulation was implemented to increase the expression of key nodes in rapamycin biosynthetic pathway, its titers may be further increased.

For each key node we selected for dynamic regulation, there exist 3-4 predicted encoding genes (isozyme genes). So far, their functions remain to be functionally identified. According to growth arrest phenotypes after static gene regulation, we could confirm that they are all required for normal cell growth and none of them are redundant genes in *S. rapamycinicus*. EQCi-mediated repression of all these genes resulted in obviously increased rapamycin production. Here, we only selected one gene from each node, whose dynamic regulation resulted in the most significantly enhanced rapamycin production, to test the effect of combinatorial repression of three essential pathways. Through screening a library of 18 engineered strains harboring different sgRNA combinations (2-3 sgRNA for each gene with different repression strength), we obtained the strain with a final rapamycin titer of 1836±191 mg/L, showing approximately 660% higher than that of the parental strain. It could be imaged that if more sgRNAs for each gene were designed and more isozyme genes were included, we may obtain strains with higher rapamycin titers by screening a large library with the help of high-throughput screening technology.

Prior to our work, static metabolic engineering has been already employed to enhance rapamycin production. For example, chromosomal integration of an additional copy of two cluster-situated activator genes *rapG* and *rapH* resulted in 20-32% and 27-55% increases in rapamycin titers (*34*). In another study, based on a genome-scale metabolic model, *pfk* (encoding 6-phosphofructokinase) was deleted, and two target genes, including *dahp* (encoding a DAHP synthase) and *rapK* (encoding a chorismatase) were overexpressed, gave approximately 30.8%, 36.2% and 44.8% increases in rapamycin titers, respectively. Combinatorial metabolic engineering by *pfk* knockout and co-overexpression of *dahP* and *rapK* led to further enhanced rapamycin production by 142.3%, achieving 250.8 mg/L rapamycin titer (*35*). Obviously, compared to previous studies with static metabolic engineering approaches, our EQCi-based regulation of multiple key nodes in essential pathways is much more effective for rapamycin titer improvement. Rapamycin, as a complex natural product of polyketide, provides a good example for EQCi system-mediated titer improvement of other polyketide compounds. Polyketide biosynthesis needs a variety of common precursors, such as malonyl-CoA, methymalonyl-CoA and isobutyryl-CoA, etc, which are mainly derived from acetyl-CoA. Therefore, essential pathways tested here, including the TCA cycle and FA synthesis pathway, can be the universal engineering targets for EQCi regulation to boost polyketide titers. Nevertheless, testing and screening of key nodes in the TCA cycle and FA pathway may be needed to identify the best targets for different specific products.

## Materials and methods

### Bacterial strains, plasmids and growth conditions

Plasmids and bacterial strains used in this study are listed in Table S1. *E. coli* strains, including DH5α and ET12567/pUZ8002, were grown at 37°C in Luria -Bertani (LB) broth or on LB agar plates. DH5α was used for general cloning and ET12567/pUZ8002 was employed as the donor strain for intergenic conjugation between *E. coli* and *Streptomyces. S. rapamycinicus* 2001 was derived from the wild-type strain NRRL 5491 by recursive physical and chemical mutagenesis, and grown at 30°C on oat agar medium (20 g/liter oat, 20 g/liter agar, pH 6.8) for preparing spore suspension and on M-ISP4 (2 g/liter tryptone, 1 g/liter yeast extract, 5 g/liter soya flour, 5 g/liter mannitol, 5 g/liter starch, 2 g/liter ammonium sulfate, 2 g/liter calcium carbonate, 1 g/liter sodium chloride, 0.5 g/liter valine, 20 g/liter agar) for intergenic conjugation. When necessary, apramycin (50 μg/mL), kanamycin (50 μg/mL) and nalidixic acid (25 μg/mL) were added.

### β-galactosidase reporter assays

The codon optimized thermophilic *lacZ* gene (*32*) under the control of the GBL-responsive promoter *srbAp* or the strong constitutive promoter and *ermEp\** (GenScript, Nanjing, China) was cloned in the integrative vector pSET152 between *Nde*I and *Xba*I to yield the plasmids of pSET-*srbAp-lacZ* and pSET-*<u>ermEp</u>*\**-lacZ*. Two plasmids were individually transferred into *S. rapamycinicus* 2001 by intergenic conjugation. Strains harboring the reporter plasmids were plated on oat agar medium and cultured at 30°C for 7 days. Strain with the empty plasmid pSET152 was used as a control. Agar cultures were inoculated into 25 ml of seed medium (15 g/liter soluble starch, 3 g/liter peptone, 0.5 g/liter L-lysine, 1.0 g/liter K_2_HPO_4_ · 3H_2_O, 3 g/liter glucose, pH 7.0) in 250 ml flasks at 28 °C for 24 h. Then, 3 ml of seed cultures were transferred into 25 ml of fermentation medium (4 g/liter soybean flour, 5 g/liter yeast extract, 2 g/liter peptone, 10 g/liter oat, 5 g/liter glycerol, 1.5 g/liter L-lysine, 40 g/liter glucose, pH 6.5) in 250 ml flasks at 28°C. After growth for 3, 5, 7 and 9 days, cultures were harvested by centrifugation at 12, 000g for 5 min. Cell pellets were dissolved in the B-PER reagent (Thermo Scientific Pierce, USA) and vortexed for 1 min. The resulting cell lysates were heat-treated at 60 °C for 30 min to remove heat unstable proteins (such as endogenous β-galactosidases), followed by centrifugation for 30 min at 12, 000g. The supernatants were used for the β-galactosidase assays as previously reported (*32*). The experiments were performed with three biological replicates.

### 2. Construction of the CRISPRi plasmids

The *ermEp*\*-driving CRISPRi plasmids were constructed based on pSET-*dcas9-actII4-NT-S1* by simply replacing the 20-nt specific guide sequence of sgRNA targeting genes of interest according to the method described previously (*36*). The EQCi plasmids pSET-*srbAp-dcas9*-*sgRNA* for dynamic regulation of single genes were generated based on the *ermEp*\*-driving CRISPRi plasmids by replacing the constitutive strong promoter *ermEp*\* with *srbAp* promoter. Briefly, the *srbAp* promoter was amplified from the genome of *S. rapamycinicus* 2001 by PCR using the primer pair *srbAp*-F/R (Table S2). The PCR product was digested with *Xba*I and *Nde*I and cloned into pSET-*ermEp*\**-dcas9*-*sgRNA*, which was treated with two same enzymes, thus generating pSET-*ermEp*\**-dcas9*-*sgRNA*. Replacement of 20 nt specific target sequences of sgRNAs in the *ermEp*\*-driving CRISPRi or the EQCi plasmids could be easily performed by the method described previously (*36*). The EQCi plasmids for multiplex gene repression (harboring multiple sgRNA cassettes) were constructed by GenScript (Nanjing, China).

### *S. rapamycinicus* fermentation and HPLC analysis of rapamycin production

Spores of *S. rapamycinicus* strains were streaked on oat agar medium and cultured at 30°C for 7 days. Agar cultures were inoculated into 25 ml of seed medium in 250 ml flasks at 28 °C for 24 h. Then, 3 ml of seed cultures were transferred into 25 ml of fermentation medium in 250 ml flasks. 0.5 ml of fermentation cultures were extracted with equal volume of methanol at room temperature for 1 h. After centrifugation at 12, 000 rpm for 10 min, supernatants were analyzed by HPLC using a reversed-phase column (Shimadzu, shim-pack VP-ODS, 4.6 um, 4.6×150mm). A mixture of water/methanol (15:85) was used as the mobile phase, with a ?ow rate of 1.0 mL/min and a retention time of 12 min. Rapamycin was monitored at a wavelength of 277.4 nm. The experiments were carried out with three biological replicates.

### Determination of *S. rapamycinicus* growth curve

The biomass of *S. rapamycinicus* strains were measured by a simplified diphenylamine colorimetric method as described before (*37*). In brief, *S. rapamycinicus* strains were grown on oat agar medium at 30°C for 7 days. Agar cultures were inoculated into 25 ml of seed medium in 250 ml flasks at 28 °C for 24 h. Subsequently, 3 ml of seed cultures were transferred into 25 ml of fermentation medium in 250 ml flasks at 28 °C. Samples (0.5 ml) were collected at an interval of 12 hours, followed by centrifugation at 12, 000g for 5 min. Cell pellets were dissolved in the diphenylamine reaction buffer (diphenylamine 1.5g, glacial acetic acid 100 ml, concentrated sulfuric acid 1.5 ml, 1.6% aqueous acetaldehyde 1 ml) and vortexed for 1 min. The reactions were incubated for 1 hour at 60°C, followed by centrifugation for 1 min at 12, 000g. OD_595_ of the supernatants were assayed by the multifunctional microtiter plate reader Tecan Spark (Tecan, Switzerland). The experiments were performed with three biological replicates.

### Quantification of mRNA

Real-time RT-PCR (RT-qPCR) analysis was conducted as reported previously (*38*). Briefly, cultures were taken at different time points (mainly at 3, 5, 7, and 9 days) and frozen quickly in liquid nitrogen and then ground into powder. RNA samples were extracted with the Ultrapure RNA Kit (SparkJade, Shanghai, China) and followed by digestion with DNase I (SparkJade, Shanghai, China) to remove residual chromosomal DNA. RT-qPCR was carried out in a MyiQ2 thermal cycler (Bio-Rad, USA), using a SYBR Green PCR premix kit (Bio-Rad, USA). PCR analysis was carried out in triplicate (technical replicates). Transcript levels of each tested gene were normalized to *hrdB* (*M271_14880*, an internal control) and the relative fold change of gene transcription was then determined by the 2^-ΔΔCT^ method (*39*). The values are averages of three independent RT-qPCR analysis. The control strain is 2001/*pdcas9* (the parental strain containing the plasmid pSET-*srbAp*-*dcas9*) or 2001/*permE*-dcas9* (the parental strain containing the plasmid pSET-*ermEp\**-*dcas9*). The experiments were conducted with three independent RNA samples (biological replicates). Error bars indicate the standard deviations (SD). The primers used are listed in Table S2.

## Acknowledgements

This work was financially supported by the National Key Research and Development Program (2019YFA0905400 and 2018YFA0903700), the National Natural Science Foundation of China (31770088 and 31630003), the National Mega-project for Innovative Drugs (2018ZX09711001-006-012) and the Funds for Creative Research Groups of China (31921006).

## Author contributions

J.T and G. Y conceived, performed experiments and drafted the manuscript. X.S provided the parental strain of *S. rapamycinicus* and medium formula for bacterial fermentations. Y.G, Y.L and W.J analyzed, interpreted the results and contributed to the writing and editing of the manuscript.

## Data and materials availability

All data needed to evaluate the conclusions in the paper are present in the paper and/or the Supplementary Materials. Additional data related to this paper may be requested from the authors.

