## Supplementary Materials for "Developing an endogenous quorum-sensing based CRISPRi circuit for autonomous and tunable dynamic regulation of multiple targets in industrial *Streptomyces*" (from another version of this preprint: version 1, 2020-01-17)

**Table S1. Strains and plasmids used in this study**

| Strains and plasmids | Genotype | References |
|---|---|---|
| **Strains** | | |
| *E. coli* strains | | |
| DH5α | F-*Φ*80 *lac*ZΔM15 Δ(*lacZYA-argF*) *U169 deoR recA1 endA1 hsdR17* s*upE44 λ*⁻ *thi-1 gyrA96 relA1* | Gicbo-BRL |
| ET12567/pUZ8002 | *dam-13*::Tn9 *dcm-6 hsdM*; harboring the non-transmissible RP4 derivative plasmid pUZ8002 | (*1*) |
| *S. rapamycinicus* 2001 | The parental strain derived from the wild-type *S. rapamycinicus* NRRL 5491 | This study |
| 2001/*sg-fabH1* | 2001 with the EQCi plasmid pSET-*srbAp-dcas9/sg-fabH1* | This study |
| 2001/*sg-fabH2* | 2001 with the EQCi plasmid pSET-*srbAp-dcas9/sg-fabH2* | This study |
| 2001/*sg-fabH3* | 2001 with the EQCi plasmid pSET-*srbAp-dcas9/sg-fabH3* | This study |
| 2001/*ermE*/sg-fabH1* | 2001 with the *ermEp**-driving CRISPRi plasmid pSET-*ermEp*-dcas9/sg-fabH1* | This study |
| 2001/*ermE*/sg-fabH2* | 2001 with the *ermEp**-driving CRISPRi plasmid pSET-*ermEp*-dcas9/sg-fabH2* | This study |
| 2001/*ermE*/sg-fabH3* | 2001 with the *ermEp**-driving CRISPRi plasmid pSET-*ermEp*-dcas9/sg-fabH3* | This study |
| 2001/*sg-gltA* | 2001 with the EQCi plasmid pSET-*srbAp-dcas9/sg-gltA* | This study |
| 2001/*sg-gltA1* | 2001 with the EQCi plasmid pSET-*srbAp-dcas9/sg-gltA1* | This study |
| 2001/*sg-gltA2* | 2001 with the EQCi plasmid pSET-*srbAp-dcas9/sg-gltA2* | This study |
| 2001/*ermE*/sg-gltA* | 2001 with the *ermEp**-driving CRISPRi plasmid pSET-*ermEp*-dcas9/sg-gltA* | This study |
| 2001/*ermE*/sg-gltA1* | 2001 with the *ermEp**-driving CRISPRi plasmid pSET-*ermEp*-dcas9/sg-gltA1* | This study |
| 2001/*ermE*/sg-gltA2* | 2001 with the *ermEp**-driving CRISPRi plasmid pSET-*ermEp*-dcas9/sg-gltA2* | This study |
| 2001/*sg-cm1* | 2001 with the EQCi plasmid pSET-*srbAp-dcas9/sg-cm1* | This study |
| 2001/*sg-cm2* | 2001 with the EQCi plasmid pSET-*srbAp-dcas9/sg-cm2* | This study |
| 2001/*sg-cm3* | 2001 with the EQCi plasmid pSET-*srbAp-dcas9/sg-cm3* | This study |
| 2001/*sg-cm4* | 2001 with the EQCi plasmid pSET-*srbAp-* | This study |

| | | |
|---|---|---|
| | *dcas9/sg-cm4* | |
| 2001/*ermE*/sg-cm1* | 2001 with the *ermEp**-driving CRISPRi plasmid pSET-*srbAp-dcas9/sg-cm1* | This study |
| 2001/*ermE*/sg-cm2* | 2001 with the *ermEp**-driving CRISPRi plasmid pSET-*srbAp-dcas9/sg-cm2* | This study |
| 2001/*ermE*/sg-cm3* | 2001 with the *ermEp**-driving CRISPRi plasmid pSET-*srbAp-dcas9/sg-cm3* | This study |
| 2001/*ermE*/sg-cm4* | 2001 with the *ermEp**-driving CRISPRi plasmid pSET-*srbAp-dcas9/sg-cm4* | This study |
| 2001/*sg1-fabH3* | 2001 with the EQCi plasmid pSET-*srbAp-dcas9/sg1-fabH3* | This study |
| 2001/*sg2-fabH3* | 2001 with the EQCi plasmid pSET-*srbAp-dcas9/sg2-fabH3* | This study |
| 2001/*sg3-fabH3* | 2001 with the EQCi plasmid pSET-*srbAp-dcas9/sg3-fabH3* | This study |
| 2001/*sg1-gltA2* | 2001 with the EQCi plasmid pSET-*srbAp-dcas9/sg1-gltA2* | This study |
| 2001/*sg2-gltA2* | 2001 with the EQCi plasmid pSET-*srbAp-dcas9/sg2-gltA2* | This study |
| 2001/*sg3-gltA2* | 2001 with the EQCi plasmid pSET-*srbAp-dcas9/sg3-gltA2* | This study |
| 2001/*sg1-cm2* | 2001 with the EQCi plasmid pSET-*srbAp-dcas9/sg1-cm2* | This study |
| 2001/*sg2-cm2* | 2001 with the EQCi plasmid pSET-*srbAp-dcas9/sg2-cm2* | This study |
| 2002 | 2001 with the EQCi plasmid pSET-*srbAp-dcas9/sg-fabH3*//sg-*gltA2/sg-cm2* | This study |
| 2003 | 2001 with the EQCi plasmid pSET-*srbAp-dcas9/sg-fabH3*/sg-*gltA2/sg2-cm2* | This study |
| 2004 | 2001 with the EQCi plasmid pSET-*srbAp-dcas9/sg-fabH3*/sg2-*gltA2/sg-cm2* | This study |
| 2005 | 2001 with the EQCi plasmid pSET-*srbAp-dcas9/sg-fabH3*/*sg2-gltA2/sg2-cm2* | This study |
| 2006 | 2001 with the EQCi plasmid pSET-*srbAp-dcas9/sg-fabH3*/sg3-*gltA2/sg-cm2* | This study |
| 2007 | 2001 with the EQCi plasmid pSET-*srbAp-dcas9/sg-fabH3/sg3-gltA2/sg2-cm2* | This study |
| 2008 | 2001 with the EQCi plasmid pSET-*srbAp-dcas9/sg2-fabH3/sg-gltA2/sg-cm2* | This study |
| 2009 | 2001 with the EQCi plasmid pSET-*srbAp-dcas9/sg2-fabH3/sg-gltA2/sg2-cm2* | This study |
| 2010 | 2001 with the EQCi plasmid pSET-*srbAp-dcas9/sg2-fabH3/sg2-gltA2/sg-cm2* | This study |
| 2011 | 2001 with the EQCi plasmid pSET-*srbAp-* | This study |

| | | |
|---|---|---|
| | *dcas9/sg2-fabH3/sg2-gltA2/sg2-cm2* | |
| 2012 | 2001 with the EQCi plasmid pSET-*srbAp-dcas9/sg2-fabH3/sg3-gltA2/sg-cm2* | This study |
| 2013 | 2001 with the EQCi plasmid pSET-*srbAp-dcas9/sg2-fabH3/sg3-gltA2/sg2-cm2* | This study |
| 2014 | 2001 with the EQCi plasmid pSET-*srbAp-dcas9/sg3-fabH3/sg-gltA2/sg-cm2* | This study |
| 2015 | 2001 with the EQCi plasmid pSET-*srbAp-dcas9/sg3-fabH3/sg-gltA2/sg2-cm2* | This study |
| 2016 | 2001 with the EQCi plasmid pSET-*srbAp-dcas9/sg3-fabH3/sg2-gltA2/sg-cm2* | This study |
| 2017 | 2001 with the EQCi plasmid pSET-*srbAp-dcas9/sg3-fabH3/sg2-gltA2/sg2-cm2* | This study |
| 2018 | 2001 with the EQCi plasmid pSET-*srbAp-dcas9/sg3-fabH3/sg3-gltA2/sg-cm2* | This study |
| 2019 | 2001 with the EQCi plasmid pSET-*srbAp-dcas9/sg3-fabH3/sg3-gltA2/sg2-cm2* | This study |
| **Plasmids** | | |
| pSET152 | An integrative plasmid, *acc(3)IV, oriT* RK2 , ΦC31 integrase/*attP* | (*2*) |
| <u>Reporter plasmids</u> | | |
| pSET-*srbAp-lacZ* | A reporter plasmid, in which the codon optimized thermophilic *lacZ* gene under the control of the GBL-responsive promoter *srbAp* was cloned between *Nde*I and *Xba*I | This study |
| pSET-*ermEp*-lacZ* | A reporter plasmid, in which the codon optimized thermophilic *lacZ* gene under the control of the constitutive strong promoter *ermEp** was cloned between *Nde*I and *Xba*I | This study |
| <u>*ermEp**-driving CRISPRi plasmids</u> | | |
| pSET-*dcas9-actII4-NT-S1* | A pSET152-derived CRISPRi plasmid, in which dCas9 is under the control of *ermEp** and the sgRNA *actII4-NT-S1* transcription cassette (targeting *actII-ORF4*, a pathway-specific activator for actinorhodin biosynthesis in *Streptomyces coelicolor*) is under the control of *j23119p* | (*3*) |
| pSET-*ermEp*-dcas9* | A control CRISPRi plasmid, in which dCas9 is under the control of *ermEp** and the sgRNA without N20 guide sequence is under the control of *j23119p* | (*3*) |
| pSET-*ermEp*-dcas9/sg-fabH1* | pSET-*ermEp*-dcas9* harboring the expression cassette of the sgRNA *sg-fabH1* | This study |

| | | |
|---|---|---|
| | targeting the nucleotide position of 8-27 on the non-template strand of *fabH1* | |
| pSET-*ermEp*-dcas9/sg-fabH2* | pSET-*ermEp*-dcas9* harboring the expression cassette of the sgRNA *sg-fabH2* targeting the nucleotide position of 27-46 on the non-template strand of *fabH2* | This study |
| pSET-*ermEp*-dcas9/sg-fabH3* | pSET-*ermEp*-dcas9* harboring the expression cassette of the sgRNA *sg-fabH3* targeting the nucleotide position of 179-198 on the non-template strand of *fabH3* | This study |
| pSET-*ermEp*-dcas9/sg-gltA* | pSET-*ermEp*-dcas9* harboring the expression cassette of the sgRNA *sg-gltA* targeting the nucleotide position of 137-156 on the non-template strand of *gltA* | This study |
| pSET-*ermEp*-dcas9/sg-gltA1* | pSET-*ermEp*-dcas9* harboring the expression cassette of the sgRNA *sg-gltA1* targeting the nucleotide position of 8-27 on the non-template strand of *gltA1* | This study |
| pSET-*ermEp*-dcas9/sg-gltA2* | pSET-*ermEp*-dcas9* harboring the expression cassette of the sgRNA *sg-gltA2* targeting the nucleotide position of 197-216 on the non-template strand of *gltA2* | This study |
| pSET-*ermEp*-dcas9/sg-cm1* | pSET-*ermEp*-dcas9* harboring the expression cassette of the sgRNA *sg-cm1* targeting the nucleotide position of 32-51 on the non-template strand of *cm1* | This study |
| pSET-*ermEp*-dcas9/sg-cm2* | pSET-*ermEp*-dcas9* harboring the expression cassette of the sgRNA *sg-cm2* targeting the nucleotide position of 149-168 on the non-template strand of *cm2* | This study |
| pSET-*ermEp*-dcas9/sg-cm3* | pSET-*ermEp*-dcas9* harboring the expression cassette of the sgRNA *sg-cm3* targeting the nucleotide position of 51-70 on the non-template strand of *cm3* | This study |
| pSET-*ermEp*-dcas9/sg-cm4* | pSET-*ermEp*-dcas9* harboring the expression cassette of the sgRNA *sg-cm4* targeting the nucleotide position of 22-41 on the non-template strand of *cm4* | This study |
| <u>EQCi plasmids</u> | | |
| pSET-*srbAp-dcas9* | A control EQCi plasmid, in which dCas9 is under the control of *srbAp* and the transcription of the sgRNA without N20 guide sequence is under the control of | This study |

| | | |
|---|---|---|
| | *j23119p* | |
| pSET-*srbAp-dcas9/sg-fabH1* | pSET-*srbAp-dcas9* harboring the expression cassette of the sgRNA *sg-fabH1* targeting the nucleotide position of 8-27 on the non-template strand of *fabH1* | This study |
| pSET-*srbAp-dcas9/sg-fabH2* | pSET-*srbAp-dcas9* harboring the expression cassette of the sgRNA *sg-fabH2* targeting the nucleotide position of 27-46 on the non-template strand of *fabH2* | This study |
| pSET-*srbAp-dcas9*/sg-*fabH3* | pSET-*srbAp-dcas9* harboring the expression cassette of the sgRNA *sg-fabH3* targeting the nucleotide position of 179-198 on the non-template strand of *fabH3* | This study |
| pSET-*srbAp-dcas9/sg-gltA* | pSET-*srbAp-dcas9* harboring the expression cassette of the sgRNA *sg-gltA* targeting the nucleotide position of 137-156 on the non-template strand of *gltA* | This study |
| pSET-*srbAp-dcas9/sg-gltA1* | pSET-*srbAp-dcas9* harboring the expression cassette of the sgRNA *sg-gltA1* targeting the nucleotide position of 8-27 on the non-template strand of *gltA1* | This study |
| pSET-*srbAp-dcas9/sg-gltA2* | pSET-*srbAp-dcas9* harboring the expression cassette of the sgRNA *sg-gltA2* targeting the nucleotide position of 197-216 on the non-template strand of *gltA2* | This study |
| pSET-*srbAp-dcas9/sg-cm1* | pSET-*srbAp-dcas9* harboring the expression cassette of the sgRNA *sg-cm1* targeting the nucleotide position of 32-51 on the non-template strand of *cm1* | This study |
| pSET-*srbAp-dcas9/sg-cm2* | pSET-*srbAp-dcas9* harboring the expression cassette of the sgRNA *sg-cm2* targeting the nucleotide position of 149-168 on the non-template strand of *cm2* | This study |
| pSET-*srbAp-dcas9/sg-cm3* | pSET-*srbAp-dcas9* harboring the expression cassette of the sgRNA *sg-cm3* targeting the nucleotide position of 51-70 on the non-template strand of *cm3* | This study |
| pSET-*srbAp-dcas9/sg-cm4* | pSET-*srbAp-dcas9* harboring the expression cassette of the sgRNA *sg-cm4* targeting the nucleotide (nt) position of 22-41 on the non-template strand of *cm4* | This study |
| pSET-*srbAp-dcas9/sg1-fabH3* | pSET-*srbAp-dcas9* harboring the expression cassette of the sgRNA *sg1-fabH3* targeting the nucleotide (nt) position of 29-48 on the | This study |

| | | |
|---|---|---|
| | non-template strand of *fabH3* | |
| pSET-*srbAp-dcas9/sg2-fabH3* | pSET-*srbAp-dcas9* harboring the expression cassette of the sgRNA *sg2-fabH3* targeting the nucleotide (nt) position of 419-438 on the non-template strand of *fabH3* | This study |
| pSET-*srbAp-dcas9/sg3-fabH3* | pSET-*srbAp-dcas9* harboring the expression cassette of the sgRNA *sg3-fabH3* targeting the nucleotide (nt) position of 665-684 on the non-template strand of *fabH3* | This study |
| pSET-*srbAp-dcas9/sg1-gltA2* | pSET-*srbAp-dcas9* harboring the expression cassette of the sgRNA *sg1-gltA2* targeting the nucleotide (nt) position of 8-27 on the non-template strand of *gltA2* | This study |
| pSET-*srbAp-dcas9/sg2-gltA2* | pSET-*srbAp-dcas9* harboring the expression cassette of the sgRNA *sg2-gltA2* targeting the nucleotide (nt) position of 383-402 on the non-template strand of *gltA2* | This study |
| pSET-*srbAp-dcas9/sg3-gltA2* | pSET-*srbAp-dcas9* harboring the expression cassette of the sgRNA *sg3-gltA2* targeting the nucleotide (nt) position of 593-612 on the non-template strand of *gltA2* | This study |
| pSET-*srbAp-dcas9/sg1-cm2* | pSET-*srbAp-dcas9* harboring the expression cassette of the sgRNA *sg1-cm1* targeting the nucleotide (nt) position of 24-43 on the non-template strand of *cm1* | This study |
| pSET-*srbAp-dcas9/sg2-cm2* | pSET-*srbAp-dcas9* harboring the expression cassette of the sgRNA *sg2-cm2* targeting the nucleotide (nt) position of 290-309 on the non-template strand of *cm2* | This study |
| pSET-*srbAp-dcas9*/*sg-fabH3//sg-gltA2/sg-cm2* | pSET-*srbAp-dcas9* harboring the expression cassettes of the sgRNA combination of *sg-fabH3*, *sg-gltA2* and *sg-cm2* | This study |
| pSET-*srbAp-dcas9/sg-fabH3/sg-gltA2/sg2-cm2* | pSET-*srbAp-dcas9* harboring the expression cassettes of the sgRNA combination of sg-*fabH3*, sg-*gltA2* and *sg2-cm2* | This study |
| pSET-*srbAp-dcas9/sg-fabH3/sg2-gltA2/sg-cm2* | pSET-*srbAp-dcas9* harboring the expression cassettes of the sgRNA combination of sg-*fabH3*, sg2-*gltA2* and *sg-cm2* | This study |
| pSET-*srbAp-dcas9/sg-fabH3/sg2-fabH3/sg2-cm2* | pSET-*srbAp-dcas9* harboring the expression cassettes of the sgRNA combination of sg-*fabH3*, sg2-*fabH3* and *sg2-cm2* | This study |
| pSET-*srbAp-dcas9/sg-fabH3/sg3-gltA2/sg-cm2* | pSET-*srbAp-dcas9* harboring the expression cassettes of the sgRNA combination of sg- | This study |

| | | |
|---|---|---|
| | *fabH3*, sg3-*gltA2* and *sg-cm2* | |
| pSET-*srbAp-dcas9/sg-fabH3/sg3-gltA2/sg2-cm2* | pSET-*srbAp-dcas9* harboring the expression cassettes of the sgRNA combination of sg-*fabH3*, *sg3-gltA2* and *sg2-cm2* | This study |
| pSET-*srbAp-dcas9/sg2-fabH3/sg-gltA2/sg-cm2* | pSET-*srbAp-dcas9* harboring the expression cassettes of the sgRNA combination of sg2-*fabH3*, *sg-gltA2* and *sg-cm2* | This study |
| pSET-*srbAp-dcas9/sg2-fabH3/sg-gltA2/sg2-cm2* | pSET-*srbAp-dcas9* harboring the expression cassettes of the sgRNA combination of sg2-*fabH3*, *sg-gltA2* and *sg2-cm2* | This study |
| pSET-*srbAp-dcas9/sg2-fabH3/sg2-gltA2/sg-cm2* | pSET-*srbAp-dcas9* harboring the expression cassettes of the sgRNA combination of sg2-*fabH3*, *sg2-gltA2* and *sg-cm2* | This study |
| pSET-*srbAp-dcas9/sg2-fabH3/sg2-gltA2/sg2-cm2* | pSET-*srbAp-dcas9* harboring the expression cassettes of the sgRNA combination of sg2-*fabH3*, *sg2-gltA2* and *sg2-cm2* | This study |
| pSET-*srbAp-dcas9/sg2-fabH3/sg3-gltA2/sg-cm2* | pSET-*srbAp-dcas9* harboring the expression cassettes of the sgRNA combination of sg2-*fabH3*, *sg3-gltA2* and *sg-cm2* | This study |
| pSET-*srbAp-dcas9/sg2-fabH3/sg3-gltA2/sg2-cm2* | pSET-*srbAp-dcas9* harboring the expression cassettes of the sgRNA combination of sg2-*fabH3*, *sg3-gltA2* and *sg2-cm2* | This study |
| pSET-*srbAp-dcas9/sg3-fabH3/sg-gltA2/sg-cm2* | pSET-*srbAp-dcas9* harboring the expression cassettes of the sgRNA combination of sg3-*fabH3*, *sg-gltA2* and *sg-cm2* | This study |
| pSET-*srbAp-dcas9/sg3-fabH3/sg-gltA2/sg2-cm2* | pSET-*srbAp-dcas9* harboring the expression cassettes of the sgRNA combination of sg3-*fabH3*, *sg-gltA2* and *sg2-cm2* | This study |
| pSET-*srbAp-dcas9/sg3*-fabH3/*sg2-gltA2/sg-cm2* | pSET-*srbAp-dcas9* harboring the expression cassettes of the sgRNA combination of sg3-fabH3, *sg2-gltA2* and *sg-cm2* | This study |
| pSET-*srbAp*-dcas9-sg3-*fabH3/sg2-gltA2/sg2-cm2* | pSET-*srbAp-dcas9* harboring the expression cassettes of the sgRNA combination of sg3-*fabH3*, *sg2-gltA2* and *sg2-cm2* | This study |
| pSET-*srbAp-dcas9/sg3-fabH3/sg3-gltA2/sg-cm2* | pSET-*srbAp-dcas9* harboring the expression cassettes of the sgRNA combination of sg3-*fabH3*, *sg3-gltA2* and *sg-cm2* | This study |
| pSET-*srbAp-dcas9/sg3-fabH3/sg3-gltA2/sg2-cm2* | pSET-*srbAp-dcas9* harboring the expression cassettes of the sgRNA combination of *sg3-fabH3*, *sg3-gltA2* and *sg2-cm2* | This study |

**Table S2. Primers used in this study**

| Oligonucleiotide | Sequence (5′-3′) |
|---|---|
| **Primers for the amplification of sgRNA expression cassettes** | |
| sg-gltA-F | TAATACTAGT**CGCGGCGGTGTTGCCGTAAC**GTTTTAGAGCTAGAA |
| sg-gltA1-F | TAATACTAGT**TTCGAGTCCGGGTACGAAGT**GTTTTAGAGCTAGAA |
| sg-gltA2-F | TAATACTAGT**GGCCTCCAGCAACCGACCCT**GTTTTAGAGCTAGAA |
| sg-fabH1-F | TAATACTAGT**GAGCGCCAGTACGTGCGAGC**GTTTTAGAGCTAGAA |
| sg-fabH2-F | TAATACTAGT**AGCCGGGCTTTTGAGCACCC**GTTTTAGAGCTAGAA |
| sg-fabH3-F | TAATACTAGT**GTGGCGCAGTGCCTTGAGAC**GTTTTAGAGCTAGAA |
| sg-cm1-F | TAATACTAGT**CGCGGTGCCGATCACATGGG**GTTTTAGAGCTAGAA |
| sg-cm2-F | TAATACTAGT**CTGGGTGCATTTGAAGCGCT**GTTTTAGAGCTAGAA |
| sg-cm3-F | TAATACTAGT**ACCTGCTCATGCATGTGCTC**GTTTTAGAGCTAGAA |
| sg-cm4-F | TAATACTAGT**ACGGCGATCAGCGCATGGCG**GTTTTAGAGCTAGAA |
| sg1-gltA2-F | TAATACTAGT**GGCCTCCAGCAACCGACCCT**GTTTTAGAGCTAGAA |
| sg2-gltA2-F | TAATACTAGT**GGCCACGGCCTCCTCCAGCC**GTTTTAGAGCTAGAA |
| sg3-gltA2-F | TAATACTAGT**CTCCGCCAGCGCCCGCTCCA**GTTTTAGAGCTAGAA |
| sg1-fabH3-F | TAATACTAGT**GTGGCGCAGTGCCTTGAGAC**GTTTTAGAGCTAGAA |
| sg2-fabH3-F | TAATACTAGT**TTCGTCCGGGAACCGGGCAC**GTTTTAGAGCTAGAA |
| sg3-fabH3-F | TAATACTAGT**GGTCTCGATGCAGCGGTGGG**GTTTTAGAGCTAGAA |
| sg1-cm2-F | TAATACTAGT**CTGGGTGCATTTGAAGCGCT**GTTTTAGAGCTAGAA |
| sg2-cm2-F | TAATACTAGT**GATGAAGTTGAGCAGCTTCT**GTTTTAGAGCTAGAA |
| NS-F | TAATACTAGTGTTTTAGAGCTAGAA |
| sgRNA-R | TTACGAATTCGGGTGTACATCCA |
| **Primers for the amplification of QS elements** | |
| scbAR-F | AATCTCGATATCTCAGTCCTTCCCGGTCGGTGCCAG |
| scbAR-R | GGATCCGGTACCTCAGCCGGAGAACGCGGGGC |
| scbAp-F | CGACTCTAGAGCCTGCCTCCTTGTTCATGT |
| scbAp-R | TGTCCATATGGGGTCCCCCCCAGGAAT |
| srbAp-F | CGACTCTAGAGTGTTCGCCGTCCTTCCCCG |
| srbAp-R | TTCTTGTCCATATGACGATCACCTTAAAATACTAATAATGTT |
| **Primers used for RT-qPCR analysis** | |
| hrdB-F1 | TTCGCGCTCGACCTCTAGTA |
| hrdB-R1 | GCACCTCTTGAACATCGGGA |
| QT gltA1-F | GACTCGAAGGAGTCGTCGC |
| QT gltA1-R | CGAAGGAGATATGGCCCACC |
| QT gltA2-F | CCACGTTCCGGGATTCGG |
| QT gltA2-R | GATGTTCATCGGCACGGGAC |
| QT gltA-F | CACAACCCCTTCGACGAGAA |
| QT gltA-R | ATATTGGCCTGCGAGGAACC |
| QT cm1-F | ATGAGCAGCAGCGTTTCGG |
| QT cm1-R | CAGCTCCAGCAGCGTCAT |
| QT cm2-F | CACAGCACGAGCGACC |
| QT cm2-R | GTTGTCGGAGCGTGCGAT |

| | |
|---|---|
| QT cm3-F | GCGGTACGAGCGGTCC |
| QT cm3-R | GACATCGGACTTGGACAGGG |
| QT cm4-F | AGGATCTTCCGCGACCAGA |
| QT cm4-R | CCACGAGTTCGCCGTTGA |
| QT fabH1-F | GGCCAATCTGCGGATCATCG |
| QT fabH1-R | GAGAGGGCGAGCGGAATG |
| QT fabH2-F | GTGCTCGTACTCGTCGCTC |
| QT fabH2-R | GTGCCGTAGTGGATGAGGTC |
| QT fabH3-F | CGTCCTCGTCGTGAGTTTCG |
| QT fabH3-R | CCTTCTGTTTGTGAGCGACC |

Note: The restriction enzyme sites are underlined. The sequences written in boldface letters indicate the specific N20 guide sequences of different genes.

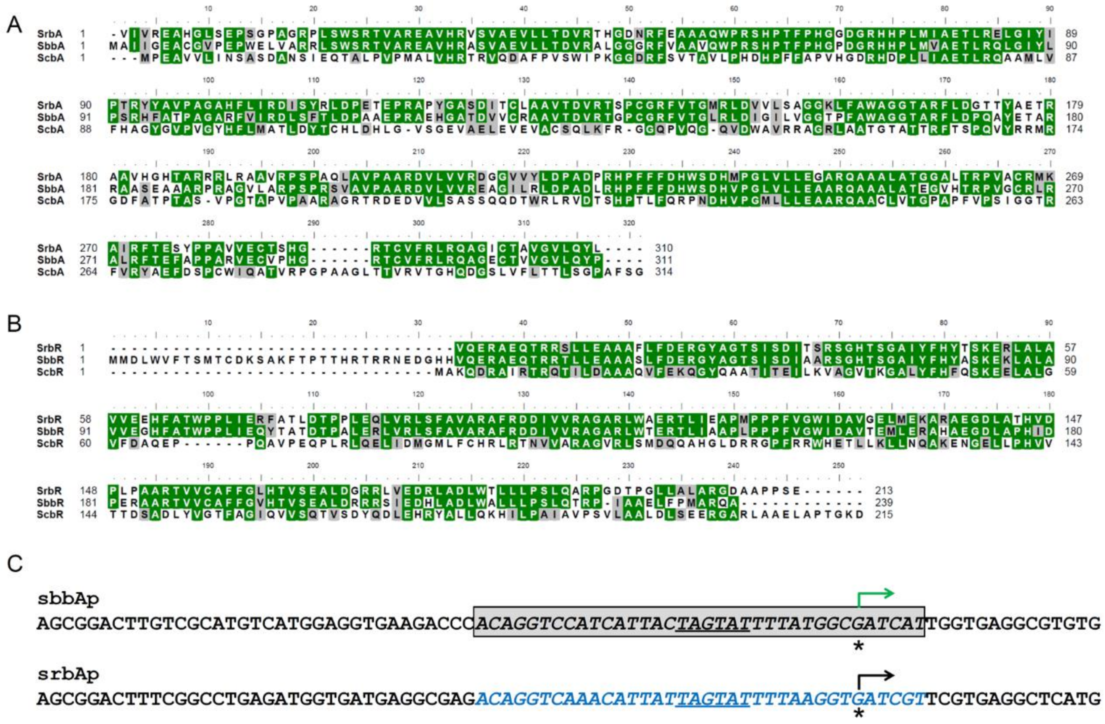

**Fig. S1 Bioinformatics analysis of the QS system in *Streptomyces rapamycinicus*.**

(A) Amino acid sequence alignment of the putative GBL synthase SrbA with other two identified GBL synthases, SbbA from *Streptomyces bingchengensis* and ScbA from *Streptomyces coelicolor*.

(B) Amino acid sequence alignment of the putative GBL receptor SrbR with other two identified GBL receptors, SbbR from *S. bingchengensis* and ScbR from *S. coelicolor*.

(C) Comparison of the promoter regions of *sbbA* from *S. bingchengensis* (*sbbAp*) and *srbA* from *S. rapamycinicus* (*srbAp*). The transcriptional start points (TSP) of *sbbAp* and *srbAp* are marked by bent arrows. The SbbR-binding sequences are boxed and the predicted SrbR-binding sequences are indicated in italicized blue letters. The putative –10 regions of *sbbAp* and *srbAp* are unlined.

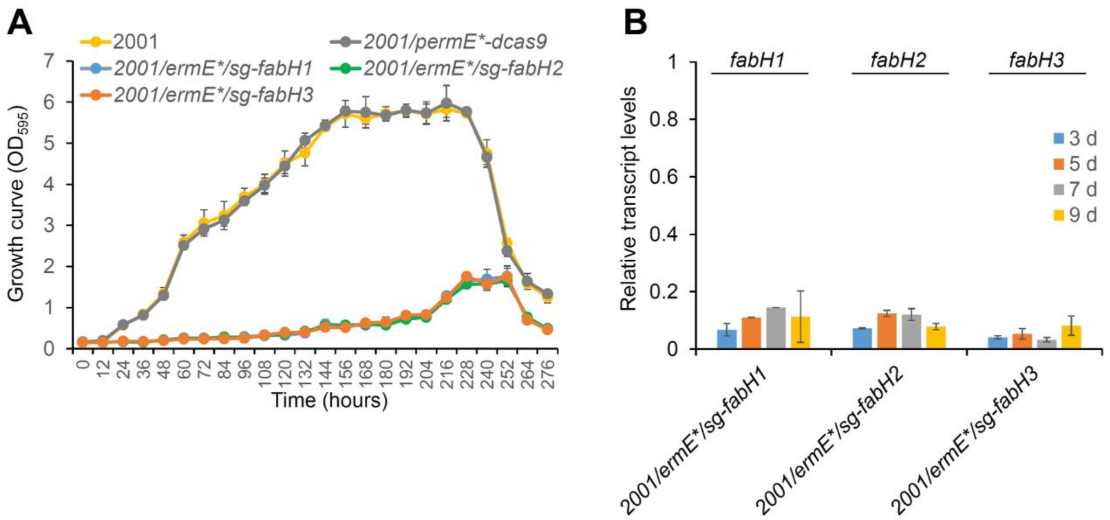


**Fig. S2 Effects of *ermEp**-driving CRISPRi circuits for static repression of the key node in FA pathway**.

(A) Growth curve of strains with *ermEp**-driving CRISPRi circuits for static repression of *fabH1-H3* (encoding FabH). For the construction of CRISPRi circuits targeting *fabH1-H3*, one sgRNA (sg) was designed for each gene. Strains with the corresponding CRISPRi circuit was indicated by 2001/*ermE*/sg-fabH1*, 2001/*ermE*/sg-fabH2* and 2001/*ermE*/sg-fabH3*, respectively. Samples were harvested from fermentation medium at the time points as indicated and the interval time is 12 hours. The parental strain 2001 and 2001 with the recombinant plasmid pSET-*ermEp*-dcas9* (2001/*permE*-dcas9*) were used as controls. Error bars denote the standard deviations (SD) of three biological replicates.

(B) Transcriptional analysis of *fabH1-H3* in strains with the corresponding *ermEp**-driving CRISPRi by RT-qPCR. RNA samples were isolated from S. *rapamycinicus* strains grown in fermentation medium at 3, 5, 7 and 9 days, respectively. *hrdB* was used as the internal control. Error bars represent the standard deviations (SD) from three biological replicates.

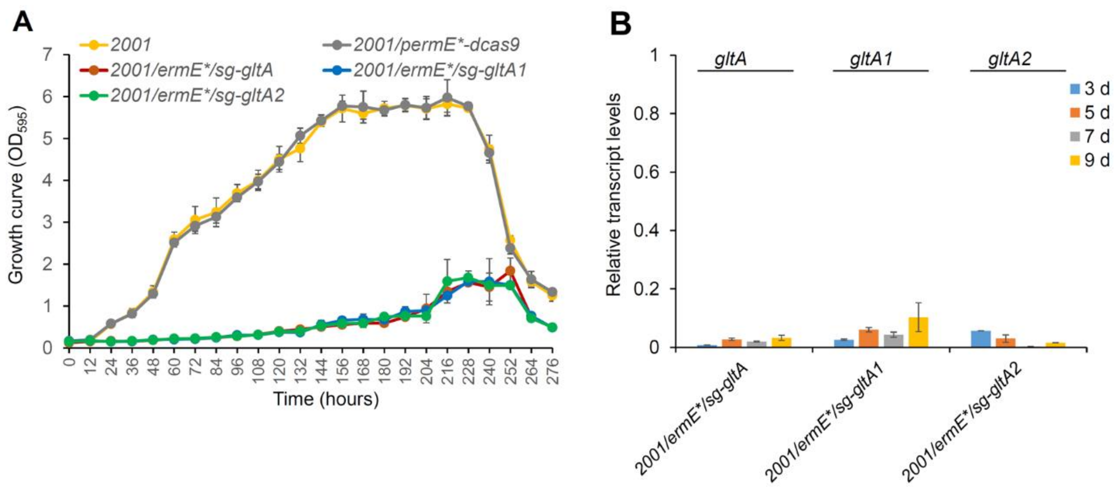


**Fig. S3 Effects of *ermEp**-driving CRISPRi circuits for static respective of the key node (citrate synthase) in TCA cycle**.

(A) Growth curve of strains with *ermEp**-driving CRISPRi circuits for static repression of *gltA-A2* (encoding citrate synthase). For the construction of CRISPRi circuits targeting *gltA-A2*, one sgRNA (sg) was designed for each gene. Strains with the corresponding CRISPRi circuit was indicated by 2001/*ermE*/sg-gltA*, 2001/*ermE*/sg-gltA1* and 2001/*ermE*/sg-gltA2*, respectively. Samples were harvested from fermentation medium at the time points as indicated and the interval time is 12 hours. The parental strain 2001 and 2001 with the recombinant plasmid pSET-*ermEp*-dcas9* (2001/*permE*-dcas9*) were used as controls. Error bars denote the standard deviations (SD) of three biological replicates

(B) Transcriptional analysis of *gltA-A2* in strains with the corresponding *ermEp**-driving CRISPRi by RT-qPCR. RNA samples were isolated from S. *rapamycinicus* strains grown in fermentation medium for 3, 5, 7 and 9 days, respectively. *hrdB* was used as the internal control. Error bars represent the standard deviations (SD) from three biological replicates.

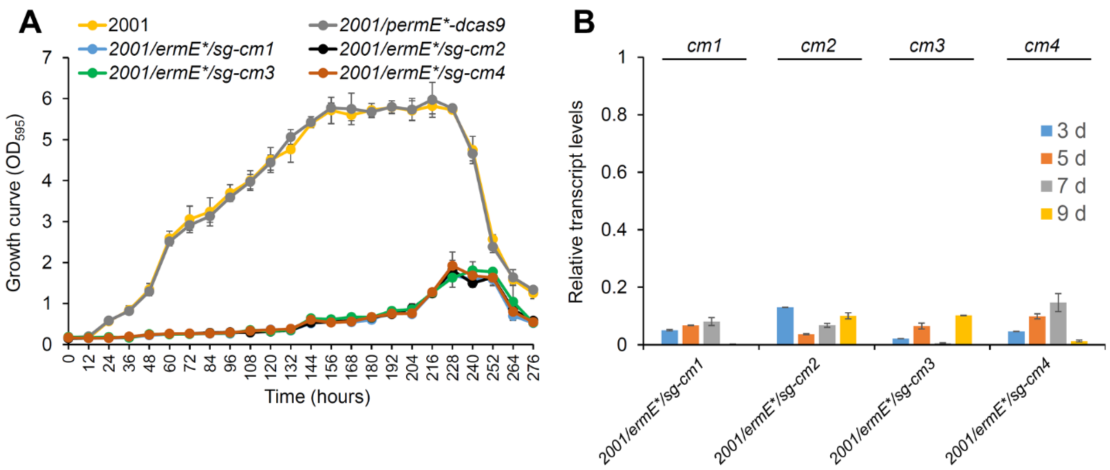


**Fig. S4 Effects of *ermEp**-driving CRISPRi circuits for static downregulation of the key node (chorismate mutase) in the AAA biosynthetic pathway**.

(A) Growth curve of strains with *ermEp**-driving CRISPRi circuits for static repression of four *cm* genes (*cm1-4*, encoding chorismate mutase). For the construction of CRISPRi circuits targeting *cm1-4*, one sgRNA (sg) was designed for each gene. Strains with the corresponding CRISPRi circuit was indicated by 2001/*ermE**/*sg-cm1*, *2001/ermE**/*sg-cm2*, 2001/*ermE**/*sg-cm3* and 2001/*ermE**/*sg-cm4*, respectively. Samples were harvested from fermentation medium at the time points as indicated and the interval time is 12 hours. The parental strain 2001 and 2001 with the recombinant plasmid pSET-*ermEp**-*dcas9* (2001/*permE**-*dcas9*) were used as controls. Error bars denote the standard deviations (SD) of three biological replicates.

(B) Transcriptional analysis of *cm1-4* in strains with the corresponding *ermEp**-driving CRISPRi by RT-qPCR. RNA samples were isolated from S. *rapamycinicus* strains grown in fermentation medium at 3, 5, 7 and 9 days, respectively. *hrdB* was used as the internal control. Error bars represent the standard deviations (SD) from three biological replicates.

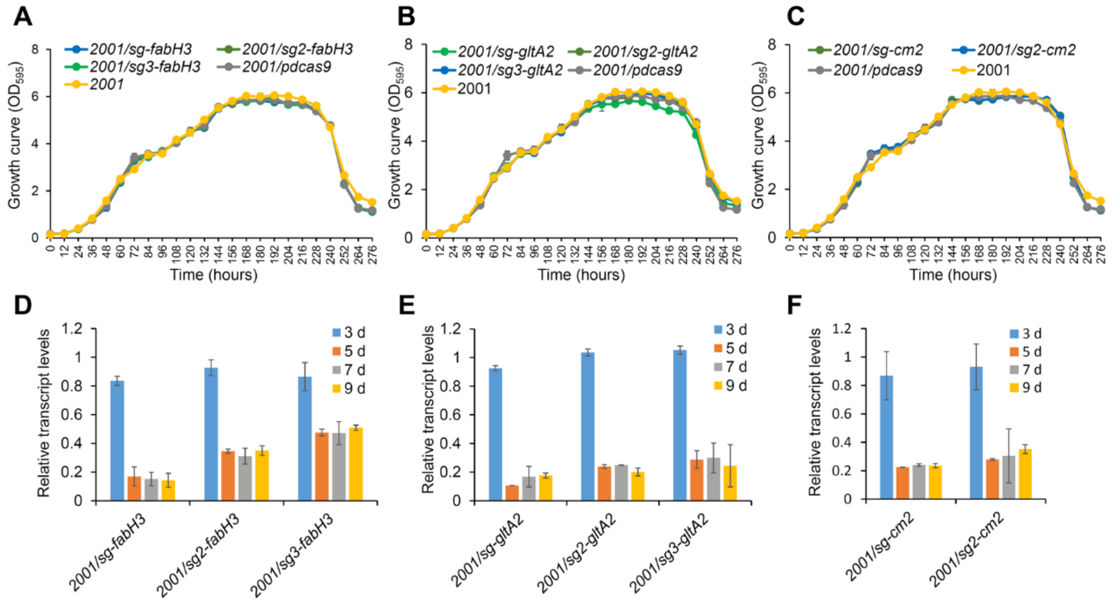


**Fig. S5 Effects of EQCi-mediated dynamic regulation with varying repression strength on cell growth and gene transcription**

(A-C) Cell growth of strains with EQCi circuits containing sgRNAs targeting different positions of *fabH3*, *gltA2* and *cm2*. Strains with the corresponding circuits were as indicated. Samples were harvested from fermentation medium at the time points as indicated and the interval time is 12 hours. The parental strain 2001 and 2001 with the recombinant plasmid pSET-*dcas9* (2001/*pdcas9*) were used as controls. Error bars denote the standard deviations (SD) of three biological replicates.

(D-F) Transcriptional level of target genes in strains with the EQCi circuits containing sgRNAs targeting different positions of *fabH3*, *gltA2* and *cm2*. RNA samples were isolated from *S. rapamycinicus* strains grown in fermentation medium for 3, 5, 7 and 9 days, respectively. *hrdB* was used as the internal control. Error bars represent the standard deviations (SD) from three biological replicates.

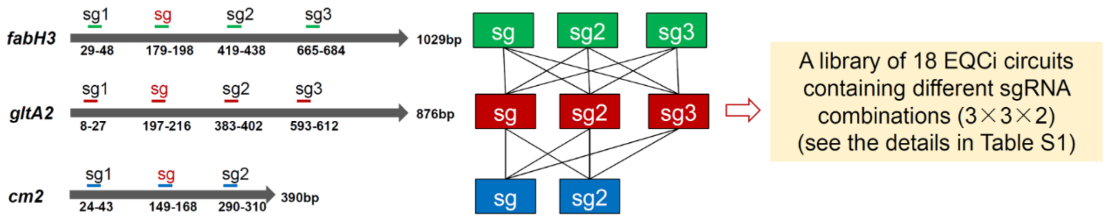


**Fig. S6 Design of a library of EQCi genetic circuits with different sgRNA combinations.**

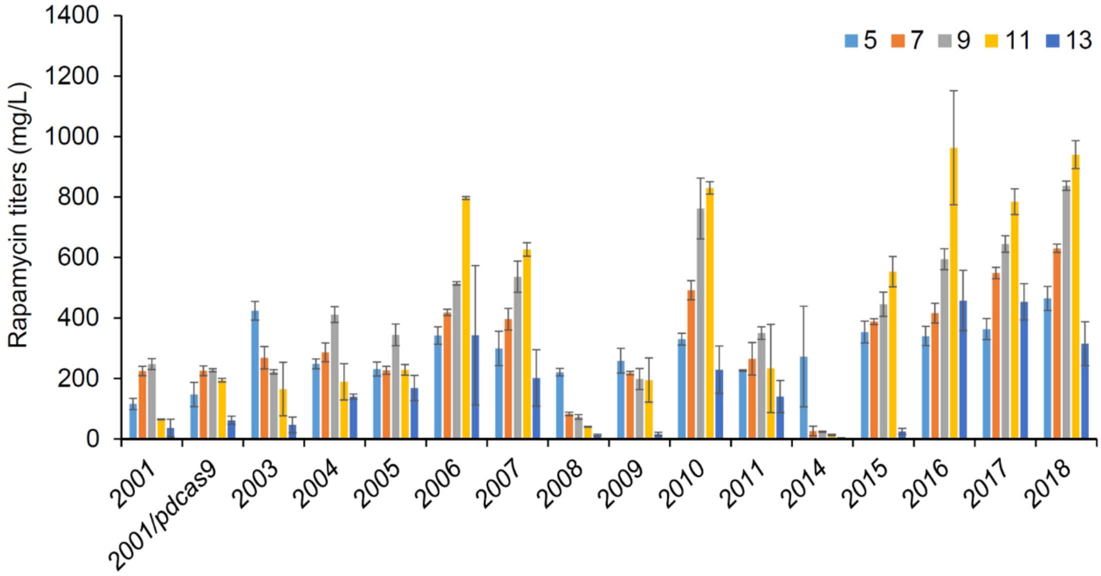


**Fig. S7 Rapamycin titers in strains with EQCi-mediated simultaneous repression of *fabH3, gltA2* and *cm2* with varying repression strength.** Tested strains were as indicated. Samples were collected from *S. rapamycinicus* strains grown in fermentation medium for 5, 7, 9, 11 and 13 days, respectively. 2001 and 2001 with the recombinant plasmid pSET-*srbAp-dcas9* (2001/*pdcas9*) were used as controls. Error bars denote the standard deviations (SD) of three biological replicates.

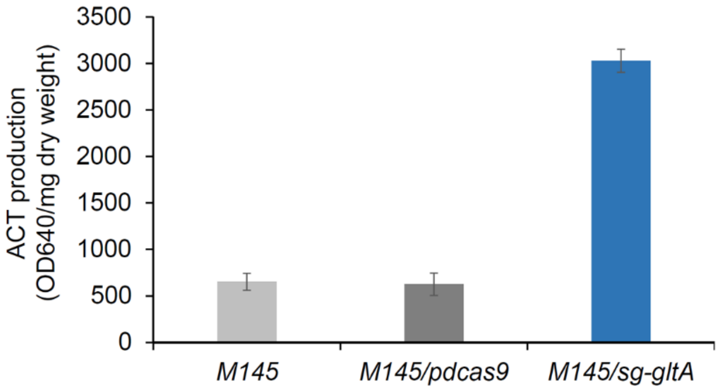


**Fig. S8 Actinorhodin (ACT) titer improvement by EQCi-mediated dynamic repression of the *gltA* gene in TCA cycle in the model strain *Streptomyces coelicolor*.**

**DNA sequences of different combinations of three sgRNA expression cassettes**.

1. *sg-fabH3*/sg-*gltA2*/*sg-cm2* (in strain 2002)

GGATCCTTGACAGCTAGCTCAGTCCTAGGTATAATACTAGT**GTGGCGCAGT**
**GCCTTGAGAC**GTTTTAGAGCTAGAAATAGCAAGTTAAAATAAGGCTAGTC
CGTTATCAACTTGAAAAAGTGGCACCGAGTCGGTGCTTTTTTTGAGTCACC
AATAAAAAACGCCCGGCGGCAACCGAGCGTTCTGAACAAATCCAGATGGA
GTTCTGAGGTCATTACTGGAGGATCCTTGACAGCTAGCTCAGTCCTAGGTAT
AATACTAGT**GGCCTCCAGCAACCGACCCT**GTTTTAGAGCTAGAAATAGCA
AGTTAAAATAAGGCTAGTCCGTTATCAACTTGAAAAAGTGGCACCGAGTCG
GTGCTTTTTTTGAGTCACCAATAAAAAACGCCCGGCGGCAACCGAGCGTTC
TGAACAAATCCAGATGGAGTTCTGAGGTCATTACTGGAGGATCCTTGACAG
CTAGCTCAGTCCTAGGTATAATACTAGT**CTGGGTGCATTTGAAGCGCT**GT
TTTAGAGCTAGAAATAGCAAGTTAAAATAAGGCTAGTCCGTTATCAACTTGA
AAAAGTGGCACCGAGTCGGTGCTTTTTTTGAGTCACCAATAAAAAACGCC
CGGCGGCAACCGAGCGTTCTGAACAAATCCAGATGGAGTTCTGAGGTCAT
TACTGGA

2. *sg-fabH3*/sg-*gltA2*/*sg2-cm2* (in strain 2003)

GGATCCTTGACAGCTAGCTCAGTCCTAGGTATAATACTAGT**GTGGCGCAGT**
**GCCTTGAGAC**GTTTTAGAGCTAGAAATAGCAAGTTAAAATAAGGCTAGTC
CGTTATCAACTTGAAAAAGTGGCACCGAGTCGGTGCTTTTTTTGAGTCACC
AATAAAAAACGCCCGGCGGCAACCGAGCGTTCTGAACAAATCCAGATGGA
GTTCTGAGGTCATTACTGGAGGATCCTTGACAGCTAGCTCAGTCCTAGGTAT
AATACTAGT**GGCCACGGCCTCCTCCAGCC**GTTTTAGAGCTAGAAATAGCA
AGTTAAAATAAGGCTAGTCCGTTATCAACTTGAAAAAGTGGCACCGAGTCG
GTGCTTTTTTTGAGTCACCAATAAAAAACGCCCGGCGGCAACCGAGCGTTC
TGAACAAATCCAGATGGAGTTCTGAGGTCATTACTGGAGGATCCTTGACAG
CTAGCTCAGTCCTAGGTATAATACTAGT**GATGAAGTTGAGCAGCTTCT**GTT
TTAGAGCTAGAAATAGCAAGTTAAAATAAGGCTAGTCCGTTATCAACTTGA
AAAAGTGGCACCGAGTCGGTGCTTTTTTTGAGTCACCAATAAAAAACGCC

CGGCGGCAACCGAGCGTTCTGAACAAATCCAGATGGAGTTCTGAGGTCATTACTGGA

3. *sg-fabH3*/sg2-*gltA2/sg-cm2* (in strain 2004)

GGATCCTTGACAGCTAGCTCAGTCCTAGGTATAATACTAGT**GTGGCGCAGTGCCTTGAGAC**GTTTTAGAGCTAGAAATAGCAAGTTAAAATAAGGCTAGTCCGTTATCAACTTGAAAAAGTGGCACCGAGTCGGTGCTTTTTTTGAGTCACCAATAAAAAACGCCCGGCGGCAACCGAGCGTTCTGAACAAATCCAGATGGAGTTCTGAGGTCATTACTGGAGGATCCTTGACAGCTAGCTCAGTCCTAGGTATAATACTAGT**GGCCACGGCCTCCTCCAGCC**GTTTTAGAGCTAGAAATAGCAAGTTAAAATAAGGCTAGTCCGTTATCAACTTGAAAAAGTGGCACCGAGTCGGTGCTTTTTTTGAGTCACCAATAAAAAACGCCCGGCGGCAACCGAGCGTTCTGAACAAATCCAGATGGAGTTCTGAGGTCATTACTGGAGGATCCTTGACAGCTAGCTCAGTCCTAGGTATAATACTAGT**CTGGGTGCATTTGAAGCGCT**GTTTTAGAGCTAGAAATAGCAAGTTAAAATAAGGCTAGTCCGTTATCAACTTGAAAAAGTGGCACCGAGTCGGTGCTTTTTTTGAGTCACCAATAAAAAACGCCCGGCGGCAACCGAGCGTTCTGAACAAATCCAGATGGAGTTCTGAGGTCATTACTGGA

4. *sg-fabH3*/sg2-*gltA2/sg2-cm2* (in strain 2005)

GGATCCTTGACAGCTAGCTCAGTCCTAGGTATAATACTAGT**GTGGCGCAGTGCCTTGAGAC**GTTTTAGAGCTAGAAATAGCAAGTTAAAATAAGGCTAGTCCGTTATCAACTTGAAAAAGTGGCACCGAGTCGGTGCTTTTTTTGAGTCACCAATAAAAAACGCCCGGCGGCAACCGAGCGTTCTGAACAAATCCAGATGGAGTTCTGAGGTCATTACTGGAGGATCCTTGACAGCTAGCTCAGTCCTAGGTATAATACTAGT**GGCCACGGCCTCCTCCAGCC**GTTTTAGAGCTAGAAATAGCAAGTTAAAATAAGGCTAGTCCGTTATCAACTTGAAAAAGTGGCACCGAGTCGGTGCTTTTTTTGAGTCACCAATAAAAAACGCCCGGCGGCAACCGAGCGTTCTGAACAAATCCAGATGGAGTTCTGAGGTCATTACTGGAGGATCCTTGACAGCTAGCTCAGTCCTAGGTATAATACTAGT**GATGAAGTTGAGCAGCTTCT**GTT

TTAGAGCTAGAAATAGCAAGTTAAAATAAGGCTAGTCCGTTATCAACTTGA
AAAAGTGGCACCGAGTCGGTGCTTTTTTTGAGTCACCAATAAAAAACGCC
CGGCGGCAACCGAGCGTTCTGAACAAATCCAGATGGAGTTCTGAGGTCAT
TACTGGA

5. *sg-fabH3*/sg3-*gltA2*/*sg-cm2* (in strain 2006)

GGATCCTTGACAGCTAGCTCAGTCCTAGGTATAATACTAGT**GTGGCGCAGT
GCCTTGAGAC**GTTTTAGAGCTAGAAATAGCAAGTTAAAATAAGGCTAGTC
CGTTATCAACTTGAAAAAGTGGCACCGAGTCGGTGCTTTTTTTGAGTCACC
AATAAAAAACGCCCGGCGGCAACCGAGCGTTCTGAACAAATCCAGATGGA
GTTCTGAGGTCATTACTGGAGGATCCTTGACAGCTAGCTCAGTCCTAGGTAT
AATACTAGT**CTCCGCCAGCGCCCGCTCCA**GTTTTAGAGCTAGAAATAGCA
AGTTAAAATAAGGCTAGTCCGTTATCAACTTGAAAAAGTGGCACCGAGTCG
GTGCTTTTTTTGAGTCACCAATAAAAAACGCCCGGCGGCAACCGAGCGTTC
TGAACAAATCCAGATGGAGTTCTGAGGTCATTACTGGAGGATCCTTGACAG
CTAGCTCAGTCCTAGGTATAATACTAGT**CTGGGTGCATTTGAAGCGCT**GT
TTTAGAGCTAGAAATAGCAAGTTAAAATAAGGCTAGTCCGTTATCAACTTGA
AAAAGTGGCACCGAGTCGGTGCTTTTTTTGAGTCACCAATAAAAAACGCC
CGGCGGCAACCGAGCGTTCTGAACAAATCCAGATGGAGTTCTGAGGTCAT
TACTGGA

6. *sg-fabH3/sg3-gltA2/sg2-cm2* (in strain 2007)

GGATCCTTGACAGCTAGCTCAGTCCTAGGTATAATACTAGT**GTGGCGCAGT
GCCTTGAGAC**GTTTTAGAGCTAGAAATAGCAAGTTAAAATAAGGCTAGTC
CGTTATCAACTTGAAAAAGTGGCACCGAGTCGGTGCTTTTTTTGAGTCACC
AATAAAAAACGCCCGGCGGCAACCGAGCGTTCTGAACAAATCCAGATGGA
GTTCTGAGGTCATTACTGGAGGATCCTTGACAGCTAGCTCAGTCCTAGGTAT
AATACTAGT**GGCCACGGCCTCCTCCAGCC**GTTTTAGAGCTAGAAATAGCA
AGTTAAAATAAGGCTAGTCCGTTATCAACTTGAAAAAGTGGCACCGAGTCG
GTGCTTTTTTTGAGTCACCAATAAAAAACGCCCGGCGGCAACCGAGCGTTC

TGAACAAATCCAGATGGAGTTCTGAGGTCATTACTGGAGGATCCTTGACAG
CTAGCTCAGTCCTAGGTATAATACTAGT**GATGAAGTTGAGCAGCTTCT**GTT
TTAGAGCTAGAAATAGCAAGTTAAAATAAGGCTAGTCCGTTATCAACTTGA
AAAAGTGGCACCGAGTCGGTGCTTTTTTTGAGTCACCAATAAAAAACGCC
CGGCGGCAACCGAGCGTTCTGAACAAATCCAGATGGAGTTCTGAGGTCAT
TACTGGA

7. *sg2-fabH3/sg-gltA2/sg-cm2* (in strain 2008)

GGATCCTTGACAGCTAGCTCAGTCCTAGGTATAATACTAGT**TTCGTCCGGG
AACCGGGCAC**GTTTTAGAGCTAGAAATAGCAAGTTAAAATAAGGCTAGTC
CGTTATCAACTTGAAAAAGTGGCACCGAGTCGGTGCTTTTTTTGAGTCACC
AATAAAAAACGCCCGGCGGCAACCGAGCGTTCTGAACAAATCCAGATGGA
GTTCTGAGGTCATTACTGGAGGATCCTTGACAGCTAGCTCAGTCCTAGGTAT
AATACTAGT**GGCCTCCAGCAACCGACCCT**GTTTTAGAGCTAGAAATAGCA
AGTTAAAATAAGGCTAGTCCGTTATCAACTTGAAAAAGTGGCACCGAGTCG
GTGCTTTTTTTGAGTCACCAATAAAAAACGCCCGGCGGCAACCGAGCGTTC
TGAACAAATCCAGATGGAGTTCTGAGGTCATTACTGGAGGATCCTTGACAG
CTAGCTCAGTCCTAGGTATAATACTAGT**CTGGGTGCATTTGAAGCGCT**GT
TTTAGAGCTAGAAATAGCAAGTTAAAATAAGGCTAGTCCGTTATCAACTTGA
AAAAGTGGCACCGAGTCGGTGCTTTTTTTGAGTCACCAATAAAAAACGCC
CGGCGGCAACCGAGCGTTCTGAACAAATCCAGATGGAGTTCTGAGGTCAT
TACTGGA

8. *sg2-fabH3/sg-gltA2/sg2-cm2* (in strain 2009)

GGATCCTTGACAGCTAGCTCAGTCCTAGGTATAATACTAGT**TTCGTCCGGG
AACCGGGCAC**GTTTTAGAGCTAGAAATAGCAAGTTAAAATAAGGCTAGTC
CGTTATCAACTTGAAAAAGTGGCACCGAGTCGGTGCTTTTTTTGAGTCACC
AATAAAAAACGCCCGGCGGCAACCGAGCGTTCTGAACAAATCCAGATGGA
GTTCTGAGGTCATTACTGGAGGATCCTTGACAGCTAGCTCAGTCCTAGGTAT
AATACTAGT**GGCCTCCAGCAACCGACCCT**GTTTTAGAGCTAGAAATAGCA

AGTTAAAATAAGGCTAGTCCGTTATCAACTTGAAAAAGTGGCACCGAGTCG
GTGCTTTTTTTGAGTCACCAATAAAAAACGCCCGGCGGCAACCGAGCGTTC
TGAACAAATCCAGATGGAGTTCTGAGGTCATTACTGGAGGATCCTTGACAG
CTAGCTCAGTCCTAGGTATAATACTAGT**GATGAAGTTGAGCAGCTTCT**GTT
TTAGAGCTAGAAATAGCAAGTTAAAATAAGGCTAGTCCGTTATCAACTTGA
AAAAGTGGCACCGAGTCGGTGCTTTTTTTGAGTCACCAATAAAAAACGCC
CGGCGGCAACCGAGCGTTCTGAACAAATCCAGATGGAGTTCTGAGGTCAT
TACTGGA

9. *sg2-fabH3/sg2-gltA2/sg-cm2* (in strain 2010)

GGATCCTTGACAGCTAGCTCAGTCCTAGGTATAATACTAGT**TTCGTCCGGG
AACCGGGCAC**GTTTTAGAGCTAGAAATAGCAAGTTAAAATAAGGCTAGTC
CGTTATCAACTTGAAAAAGTGGCACCGAGTCGGTGCTTTTTTTGAGTCACC
AATAAAAAACGCCCGGCGGCAACCGAGCGTTCTGAACAAATCCAGATGGA
GTTCTGAGGTCATTACTGGAGGATCCTTGACAGCTAGCTCAGTCCTAGGTAT
AATACTAGT**GGCCACGGCCTCCTCCAGCC**GTTTTAGAGCTAGAAATAGCA
AGTTAAAATAAGGCTAGTCCGTTATCAACTTGAAAAAGTGGCACCGAGTCG
GTGCTTTTTTTGAGTCACCAATAAAAAACGCCCGGCGGCAACCGAGCGTTC
TGAACAAATCCAGATGGAGTTCTGAGGTCATTACTGGAGGATCCTTGACAG
CTAGCTCAGTCCTAGGTATAATACTAGT**CTGGGTGCATTTGAAGCGCT**GT
TTTAGAGCTAGAAATAGCAAGTTAAAATAAGGCTAGTCCGTTATCAACTTGA
AAAAGTGGCACCGAGTCGGTGCTTTTTTTGAGTCACCAATAAAAAACGCC
CGGCGGCAACCGAGCGTTCTGAACAAATCCAGATGGAGTTCTGAGGTCAT
TACTGGA

10. *sg2-fabH3/sg2-gltA2/sg2-cm2* (in strain 2011)

GGATCCTTGACAGCTAGCTCAGTCCTAGGTATAATACTAGT**TTCGTCCGGG
AACCGGGCAC**GTTTTAGAGCTAGAAATAGCAAGTTAAAATAAGGCTAGTC
CGTTATCAACTTGAAAAAGTGGCACCGAGTCGGTGCTTTTTTTGAGTCACC
AATAAAAAACGCCCGGCGGCAACCGAGCGTTCTGAACAAATCCAGATGGA

GTTCTGAGGTCATTACTGGAGGATCCTTGACAGCTAGCTCAGTCCTAGGTAT
AATACTAGT**GGCCACGGCCTCCTCCAGCC**GTTTTAGAGCTAGAAATAGCA
AGTTAAAATAAGGCTAGTCCGTTATCAACTTGAAAAAGTGGCACCGAGTCG
GTGCTTTTTTTGAGTCACCAATAAAAAACGCCCGGCGGCAACCGAGCGTTC
TGAACAAATCCAGATGGAGTTCTGAGGTCATTACTGGAGGATCCTTGACAG
CTAGCTCAGTCCTAGGTATAATACTAGT**GATGAAGTTGAGCAGCTTCT**GTT
TTAGAGCTAGAAATAGCAAGTTAAAATAAGGCTAGTCCGTTATCAACTTGA
AAAAGTGGCACCGAGTCGGTGCTTTTTTTGAGTCACCAATAAAAAACGCC
CGGCGGCAACCGAGCGTTCTGAACAAATCCAGATGGAGTTCTGAGGTCAT
TACTGGA

11. *sg2-fabH3/sg3-gltA2/sg-cm2* (in strain 2012)
GGATCCTTGACAGCTAGCTCAGTCCTAGGTATAATACTAGT**TTCGTCCGGG
AACCGGGCAC**GTTTTAGAGCTAGAAATAGCAAGTTAAAATAAGGCTAGTC
CGTTATCAACTTGAAAAAGTGGCACCGAGTCGGTGCTTTTTTTGAGTCACC
AATAAAAAACGCCCGGCGGCAACCGAGCGTTCTGAACAAATCCAGATGGA
GTTCTGAGGTCATTACTGGAGGATCCTTGACAGCTAGCTCAGTCCTAGGTAT
AATACTAGT**CTCCGCCAGCGCCCGCTCCA**GTTTTAGAGCTAGAAATAGCA
AGTTAAAATAAGGCTAGTCCGTTATCAACTTGAAAAAGTGGCACCGAGTCG
GTGCTTTTTTTGAGTCACCAATAAAAAACGCCCGGCGGCAACCGAGCGTTC
TGAACAAATCCAGATGGAGTTCTGAGGTCATTACTGGAGGATCCTTGACAG
CTAGCTCAGTCCTAGGTATAATACTAGT**CTGGGTGCATTTGAAGCGCT**GT
TTTAGAGCTAGAAATAGCAAGTTAAAATAAGGCTAGTCCGTTATCAACTTGA
AAAAGTGGCACCGAGTCGGTGCTTTTTTTGAGTCACCAATAAAAAACGCC
CGGCGGCAACCGAGCGTTCTGAACAAATCCAGATGGAGTTCTGAGGTCAT
TACTGGA

12. *sg2-fabH3/sg3-gltA2/sg2-cm2* (in strain 2013)
GGATCCTTGACAGCTAGCTCAGTCCTAGGTATAATACTAGT**TTCGTCCGGG
AACCGGGCAC**GTTTTAGAGCTAGAAATAGCAAGTTAAAATAAGGCTAGTC

CGTTATCAACTTGAAAAAGTGGCACCGAGTCGGTGCTTTTTTTGAGTCACC
AATAAAAAACGCCCGGCGGCAACCGAGCGTTCTGAACAAATCCAGATGGA
GTTCTGAGGTCATTACTGGAGGATCCTTGACAGCTAGCTCAGTCCTAGGTAT
AATACTAGT**CTCCGCCAGCGCCCGCTCCA**GTTTTAGAGCTAGAAATAGCA
AGTTAAAATAAGGCTAGTCCGTTATCAACTTGAAAAAGTGGCACCGAGTCG
GTGCTTTTTTTGAGTCACCAATAAAAAACGCCCGGCGGCAACCGAGCGTTC
TGAACAAATCCAGATGGAGTTCTGAGGTCATTACTGGAGGATCCTTGACAG
CTAGCTCAGTCCTAGGTATAATACTAGT**GATGAAGTTGAGCAGCTTCT**GTT
TTAGAGCTAGAAATAGCAAGTTAAAATAAGGCTAGTCCGTTATCAACTTGA
AAAGTGGCACCGAGTCGGTGCTTTTTTTGAGTCACCAATAAAAAACGCC
CGGCGGCAACCGAGCGTTCTGAACAAATCCAGATGGAGTTCTGAGGTCAT
TACTGGA

13. *sg3-fabH3/sg-gltA2/sg-cm2* (in strain 2014)

GGATCCTTGACAGCTAGCTCAGTCCTAGGTATAATACTAGT**GGTCTCGATG
CAGCGGTGGG**GTTTTAGAGCTAGAAATAGCAAGTTAAAATAAGGCTAGTC
CGTTATCAACTTGAAAAAGTGGCACCGAGTCGGTGCTTTTTTTGAGTCACC
AATAAAAAACGCCCGGCGGCAACCGAGCGTTCTGAACAAATCCAGATGGA
GTTCTGAGGTCATTACTGGAGGATCCTTGACAGCTAGCTCAGTCCTAGGTAT
AATACTAGT**GGCCTCCAGCAACCGACCCT**GTTTTAGAGCTAGAAATAGCA
AGTTAAAATAAGGCTAGTCCGTTATCAACTTGAAAAAGTGGCACCGAGTCG
GTGCTTTTTTTGAGTCACCAATAAAAAACGCCCGGCGGCAACCGAGCGTTC
TGAACAAATCCAGATGGAGTTCTGAGGTCATTACTGGAGGATCCTTGACAG
CTAGCTCAGTCCTAGGTATAATACTAGT**CTGGGTGCATTTGAAGCGCT**GT
TTTAGAGCTAGAAATAGCAAGTTAAAATAAGGCTAGTCCGTTATCAACTTGA
AAAGTGGCACCGAGTCGGTGCTTTTTTTGAGTCACCAATAAAAAACGCC
CGGCGGCAACCGAGCGTTCTGAACAAATCCAGATGGAGTTCTGAGGTCAT
TACTGGA

14. *sg3-fabH3/sg-gltA2/sg2-cm2* (in strain 2015)

GGATCCTTGACAGCTAGCTCAGTCCTAGGTATAATACTAGT**GGTCTCGATG**
**CAGCGGTGGG**GTTTTAGAGCTAGAAATAGCAAGTTAAAATAAGGCTAGTC
CGTTATCAACTTGAAAAAGTGGCACCGAGTCGGTGCTTTTTTTGAGTCACC
AATAAAAAACGCCCGGCGGCAACCGAGCGTTCTGAACAAATCCAGATGGA
GTTCTGAGGTCATTACTGGAGGATCCTTGACAGCTAGCTCAGTCCTAGGTAT
AATACTAGT**GGCCTCCAGCAACCGACCCT**GTTTTAGAGCTAGAAATAGCA
AGTTAAAATAAGGCTAGTCCGTTATCAACTTGAAAAAGTGGCACCGAGTCG
GTGCTTTTTTTGAGTCACCAATAAAAAACGCCCGGCGGCAACCGAGCGTTC
TGAACAAATCCAGATGGAGTTCTGAGGTCATTACTGGAGGATCCTTGACAG
CTAGCTCAGTCCTAGGTATAATACTAGT**GATGAAGTTGAGCAGCTTCT**GTT
TTAGAGCTAGAAATAGCAAGTTAAAATAAGGCTAGTCCGTTATCAACTTGA
AAAAGTGGCACCGAGTCGGTGCTTTTTTTGAGTCACCAATAAAAAACGCC
CGGCGGCAACCGAGCGTTCTGAACAAATCCAGATGGAGTTCTGAGGTCAT
TACTGGA

15. *sg3-fabH3/sg2-gltA2/sg-cm2* (in strain 2016)

GGATCCTTGACAGCTAGCTCAGTCCTAGGTATAATACTAGT**GGTCTCGATG**
**CAGCGGTGGG**GTTTTAGAGCTAGAAATAGCAAGTTAAAATAAGGCTAGTC
CGTTATCAACTTGAAAAAGTGGCACCGAGTCGGTGCTTTTTTTGAGTCACC
AATAAAAAACGCCCGGCGGCAACCGAGCGTTCTGAACAAATCCAGATGGA
GTTCTGAGGTCATTACTGGAGGATCCTTGACAGCTAGCTCAGTCCTAGGTAT
AATACTAGT**GGCCACGGCCTCCTCCAGCC**GTTTTAGAGCTAGAAATAGCA
AGTTAAAATAAGGCTAGTCCGTTATCAACTTGAAAAAGTGGCACCGAGTCG
GTGCTTTTTTTGAGTCACCAATAAAAAACGCCCGGCGGCAACCGAGCGTTC
TGAACAAATCCAGATGGAGTTCTGAGGTCATTACTGGAGGATCCTTGACAG
CTAGCTCAGTCCTAGGTATAATACTAGT**CTGGGTGCATTTGAAGCGCT**GT
TTTAGAGCTAGAAATAGCAAGTTAAAATAAGGCTAGTCCGTTATCAACTTGA
AAAAGTGGCACCGAGTCGGTGCTTTTTTTGAGTCACCAATAAAAAACGCC
CGGCGGCAACCGAGCGTTCTGAACAAATCCAGATGGAGTTCTGAGGTCAT
TACTGGA

16. *sg3-fabH3/sg2-gltA2/sg2-cm2* (in strain 2017)

GGATCCTTGACAGCTAGCTCAGTCCTAGGTATAATACTAGT**GGTCTCGATG
CAGCGGTGGG**GTTTTAGAGCTAGAAATAGCAAGTTAAAATAAGGCTAGTC
CGTTATCAACTTGAAAAAGTGGCACCGAGTCGGTGCTTTTTTTGAGTCACC
AATAAAAAACGCCCGGCGGCAACCGAGCGTTCTGAACAAATCCAGATGGA
GTTCTGAGGTCATTACTGGAGGATCCTTGACAGCTAGCTCAGTCCTAGGTAT
AATACTAGT**GGCCACGGCCTCCTCCAGCC**GTTTTAGAGCTAGAAATAGCA
AGTTAAAATAAGGCTAGTCCGTTATCAACTTGAAAAAGTGGCACCGAGTCG
GTGCTTTTTTTGAGTCACCAATAAAAAACGCCCGGCGGCAACCGAGCGTTC
TGAACAAATCCAGATGGAGTTCTGAGGTCATTACTGGAGGATCCTTGACAG
CTAGCTCAGTCCTAGGTATAATACTAGT**GATGAAGTTGAGCAGCTTCT**GTT
TTAGAGCTAGAAATAGCAAGTTAAAATAAGGCTAGTCCGTTATCAACTTGA
AAAAGTGGCACCGAGTCGGTGCTTTTTTTGAGTCACCAATAAAAAACGCC
CGGCGGCAACCGAGCGTTCTGAACAAATCCAGATGGAGTTCTGAGGTCAT
TACTGGA

17. *sg3-fabH3/sg3-gltA2/sg-cm2* (in strain 2018)

GGATCCTTGACAGCTAGCTCAGTCCTAGGTATAATACTAGT**GGTCTCGATG
CAGCGGTGGG**GTTTTAGAGCTAGAAATAGCAAGTTAAAATAAGGCTAGTC
CGTTATCAACTTGAAAAAGTGGCACCGAGTCGGTGCTTTTTTTGAGTCACC
AATAAAAAACGCCCGGCGGCAACCGAGCGTTCTGAACAAATCCAGATGGA
GTTCTGAGGTCATTACTGGAGGATCCTTGACAGCTAGCTCAGTCCTAGGTAT
AATACTAGT**CTCCGCCAGCGCCCGCTCCA**GTTTTAGAGCTAGAAATAGCA
AGTTAAAATAAGGCTAGTCCGTTATCAACTTGAAAAAGTGGCACCGAGTCG
GTGCTTTTTTTGAGTCACCAATAAAAAACGCCCGGCGGCAACCGAGCGTTC
TGAACAAATCCAGATGGAGTTCTGAGGTCATTACTGGAGGATCCTTGACAG
CTAGCTCAGTCCTAGGTATAATACTAGT**CTGGGTGCATTTGAAGCGCT**GT
TTTAGAGCTAGAAATAGCAAGTTAAAATAAGGCTAGTCCGTTATCAACTTGA
AAAAGTGGCACCGAGTCGGTGCTTTTTTTGAGTCACCAATAAAAAACGCC

CGGCGGCAACCGAGCGTTCTGAACAAATCCAGATGGAGTTCTGAGGTCAT
TACTGGA

18. *sg3-fabH3/sg3-gltA2/sg2-cm2* (in strain 2019)

GGATCCTTGACAGCTAGCTCAGTCCTAGGTATAATACTAGT**GGTCTCGATG
CAGCGGTGGG**GTTTTAGAGCTAGAAATAGCAAGTTAAAATAAGGCTAGTC
CGTTATCAACTTGAAAAAGTGGCACCGAGTCGGTGCTTTTTTTGAGTCACC
AATAAAAAACGCCCGGCGGCAACCGAGCGTTCTGAACAAATCCAGATGGA
GTTCTGAGGTCATTACTGGAGGATCCTTGACAGCTAGCTCAGTCCTAGGTAT
AATACTAGT**CTCCGCCAGCGCCCGCTCCA**GTTTTAGAGCTAGAAATAGCA
AGTTAAAATAAGGCTAGTCCGTTATCAACTTGAAAAAGTGGCACCGAGTCG
GTGCTTTTTTTGAGTCACCAATAAAAAACGCCCGGCGGCAACCGAGCGTTC
TGAACAAATCCAGATGGAGTTCTGAGGTCATTACTGGAGGATCCTTGACAG
CTAGCTCAGTCCTAGGTATAATACTAGT**GATGAAGTTGAGCAGCTTCT**GTT
TTAGAGCTAGAAATAGCAAGTTAAAATAAGGCTAGTCCGTTATCAACTTGA
AAAAGTGGCACCGAGTCGGTGCTTTTTTTGAGTCACCAATAAAAAACGCC
CGGCGGCAACCGAGCGTTCTGAACAAATCCAGATGGAGTTCTGAGGTCAT
TACTGGA

Note:

Green letters: the *j23119* promoter. Red letters: the specific N20 guide sequences of different genes. Black letters: sgRNA scaffold. Grey letters: T0 terminators.


1. B. Gust, G. L. Challis, K. Fowler, T. Kieser, K. F. Chater, PCR-targeted Streptomyces gene replacement identifies a protein domain needed for biosynthesis of the sesquiterpene soil odor geosmin. *Proc Natl Acad Sci U S A* **100**, 1541-1546 (2003).
2. T. Kieser, M. J. Bibb, M. J. Buttner, K. F. Chater., Practical Streptomyces genetics. . *John Innes Foundation, Norwich, England.*, (2000).
3. Y. Zhao *et al.*, CRISPR/dCas9-Mediated Multiplex Gene Repression in Streptomyces. *Biotechnol J*, (2018).
